# A Cautionary Note on Using STRUCTURE to Detect Hybridization in a Phylogenetic Context

**DOI:** 10.1101/2024.02.06.579057

**Authors:** Xiao-Xu Pang, Da-Yong Zhang

## Abstract

Population genetic clustering methods are widely used to detect hybridization events between closely related populations within species, as well as between deeply diverged lineages across phylogenetic time-scales, although their strengths and limitations in the latter cases remain poorly explored. This study presents the first systematic evaluation of the performance of the most popular population clustering method, STRUCTURE, under a variety of cross-species hybridization scenarios, including hybrid speciation, as well as introgression involving ghost (i.e., extinct or unsampled) lineages or otherwise. Our simulations demonstrate that STRUCTURE performs well in identifying hybrids and their parental donors only when admixture happens very recently between sampled extant lineages. However, STRUCTURE generally fails to detect signals of admixture when hybridization occurs in deep time or when gene flow stems from ghost lineages. We find that symmetrical parental contribution in cases of hybrid speciation will often be revealed as extremely asymmetrical in STRUCTURE, especially when the admixture event occurred more than some time ago. Our results suggest that population-genetic clustering methods may be very inefficient for detecting either ancient or ghost admixtures, partly explaining why ghost introgression has escaped the attention of evolutionary biologists until recently.

## Introduction

Hybridization is a process wherein genetically distinct yet not entirely isolated lineages mate and give rise to offspring. Recent phylogenomic studies have revealed that interspecific hybridization is much more prevalent than previously recognized across the entire tree of life (Mallet 2005; Arnold 2006; Schwenk et al. 2008; Whitney et al. 2010). At times, hybridization can lead to the formation of new lineages that combine alleles from parental gene pools, generally known as hybrid speciation (Mallet 2007). More frequently, hybridization results in the transfer of genetic diversity between species through the subsequent backcrossing of hybrid offspring with their parent species, a phenomenon called introgression (Anderson 1953). Interspecific hybridization has long been a focus of inquiry in evolutionary biology, due to its widely perceived importance in speciation, adaptation to new environments, and evolutionary innovation (Mallet et al. 2016; Taylor and Larson 2019; Edelman and Mallet 2021).

Hybridization and introgression, as important as they may be, are nonetheless difficult to detect and characterize. The widespread availability of genome-scale data has spurred advances in computational and statistical methods for hybridization detection. These methods range in complexity from simple and fast summary tests like the *D*-statistic (Green et al. 2010) to model-based likelihood methods that directly reconstruct phylogenetic networks for a set of species in question (see Blair and Ané 2020; Hibbins and Hahn 2022 for reviews). These approaches typically use data from only one sample per species, even when multiple samples are available (Hibbins and Hahn 2022). It is noteworthy that summary methods often encounter non-identifiability issues, and current network inference techniques face scalability challenges, struggling with even a moderate number of taxa or sequences (Degnan 2018; Wen et al. 2018; Pang and Zhang 2024). An alternative, yet complementary, strategy involves using population genetic clustering methods to identify hybrid individuals across multiple samples per species (Pritchard et al. 2000; Alexander et al. 2009; Cheng et al. 2017), offering another window onto hybridization/introgression (Blair and Ané 2020; Stull et al. 2023). Originally designed for identifying population structure in contemporary populations, these population clustering techniques are increasingly applied in studies investigating interspecific hybridization and introgression on deep time-scales (Ru et al. 2018; White et al. 2018; Zhang et al. 2019; Barth et al. 2020; Li et al. 2020; Sun et al. 2020; Li X. et al. 2021; Dittberner et al. 2022; Zou et al. 2022; Lopes et al. 2023; Sørensen et al. 2023; Wu et al. 2023; Yu et al. 2023).

The effectiveness of population clustering methods in detecting hybridization between closely related populations that have diverged for only a short time (say, tens or hundreds of generations) has been well-documented (Bohling et al. 2013; Neophytou 2014; Oliveira et al. 2015; van Wyk et al. 2017; Toyama et al. 2020; Ravagni et al. 2021). Nonetheless, there is a doubt about whether these methods are equally adept at studying hybridization between lineages that have diverged over much longer evolutionary time-scales (say, tens of thousands of generations or longer). For example, a population-level study by Toyama et al. (2020) demonstrated that the most popular Bayesian clustering method, STRUCTURE (Pritchard et al. 2000), is sensitive to post-admixture genetic drift, which could skew the accurate detection and quantification of hybridization events. It is plausible that genetic drift, which naturally intensifies over time, would have a more pronounced impact on the detection of hybridization on deeper evolutionary scales. Simulations conducted by Kong and Kubatko (2021) confirmed certain limitations of population clustering methods when applied to phylogenetic time-scales. Their findings revealed that both STRUCTURE and ADMIXTURE (another popular clustering method implemented within a maximum likelihood framework; Alexander et al. 2009) tend to perform poorly when dealing with scenarios involving asymmetrical parental contributions to hybrids or multiple hybridization events, with ADMIXTURE being more adversely affected. Moreover, they found that an excessive amount of incomplete lineage sorting (ILS) or a large genetic divergence between hybridizing species can induce spurious admixture signals in STRUCTURE analysis. However, it is important to note that their study has a relatively narrow focus. They primarily concentrate on hybrid speciation—which might be less common in nature (Schumer et al. 2014; Taylor and Larson 2019)—and one specific form of introgression, namely, inflow introgression, which occurs in the direction from the early-diverged species *C* to one of the recently-diverged sister species *A* or *B* within the species tree *AB|C*. But other forms of introgression such as outflow introgression (from recently-diverged to early-diverged species), introgression between sister species, or ghost introgression (explained below) remain unexplored in their research.

Evolutionary studies are often constrained to a limited subset of species due to many lineages becoming either extinct or unsampled because of technical constraints or their irrelevance to the research question (Ottenburghs 2020; Tricou et al. 2022a, 2022b). The phenomenon of ‘ghost introgression’, which refers to introgression from extinct or unsampled lineages to the sampled species, is widely acknowledged and has been uncovered in a growing number of plants (e.g., Ru et al. 2018; Li M. et al. 2021; Ding et al. 2022; Li et al. 2022; Chang et al. 2023; Tiley et al. 2023) and animals (Sankararaman et al. 2014; Ai et al. 2015; Kuhlwilm et al. 2019; Wang et al. 2020; Rocha et al. 2022; Pawar et al. 2023; Yamahira et al. 2023; Kato et al. 2024). Intuitively, the involvement of ghost lineages in a gene flow event inevitably leads to a loss of information for hybridization detection, thereby negatively affecting admixture inference. For example, Lawson et al. (2018) found that relatives of extinct parental lineages might be incorrectly inferred as participating in gene flow. Yet, comprehensive and detailed research into the impact of ghost lineages on the effectiveness of population clustering methods for detecting hybridization remains poorly explored, if not completely lacking.

In this article, our goal is to examine the performance of population clustering methods under various scenarios of hybridization/introgression on phylogenetic time-scales, with a special focus on ghost introgression, which has received little attention in previous benchmark studies. To conduct our analysis, we employed the gold standard method, STRUCTURE, as a representative of such approaches. In our simulations, we set the divergence times for two lineages engaged in hybridization to be at least 1.5 coalescent units, which amount to 1.5 million years if assuming an effective population size of 10,000 and a generation length of 50 years as is often the case with long-lived temperate trees (Bai et al. 2018). We simulated SNP data under scenarios involving cross-species gene flow, including hybrid speciation (Fig. 1a), as well as various scenarios of introgression within a designated species tree *AB|C* (Figs. 2a and 3a). For the sake of clarity, we introduce a new term— horizontal introgression—to represent those introgression events where the donor lineages were present in the sample, as graphically depicted by a horizontal reticulation edge on the species tree. Conceptually, horizontal introgression serves as the counterpart of ghost introgression, which corresponds to a non-horizontal reticulation edge. Our evaluation focuses on two key aspects of STRUCTURE’s performance: (i) the identification of hybrid lineages and (ii) the estimation of contribution proportions from donors. Our findings shed light on the strengths and limitations of population clustering methods for detecting hybridization across various hybridization scenarios, providing valuable insights for biologists in the appropriate application and interpretation of these methods for evolutionary analysis.

**FIGURE 1.**
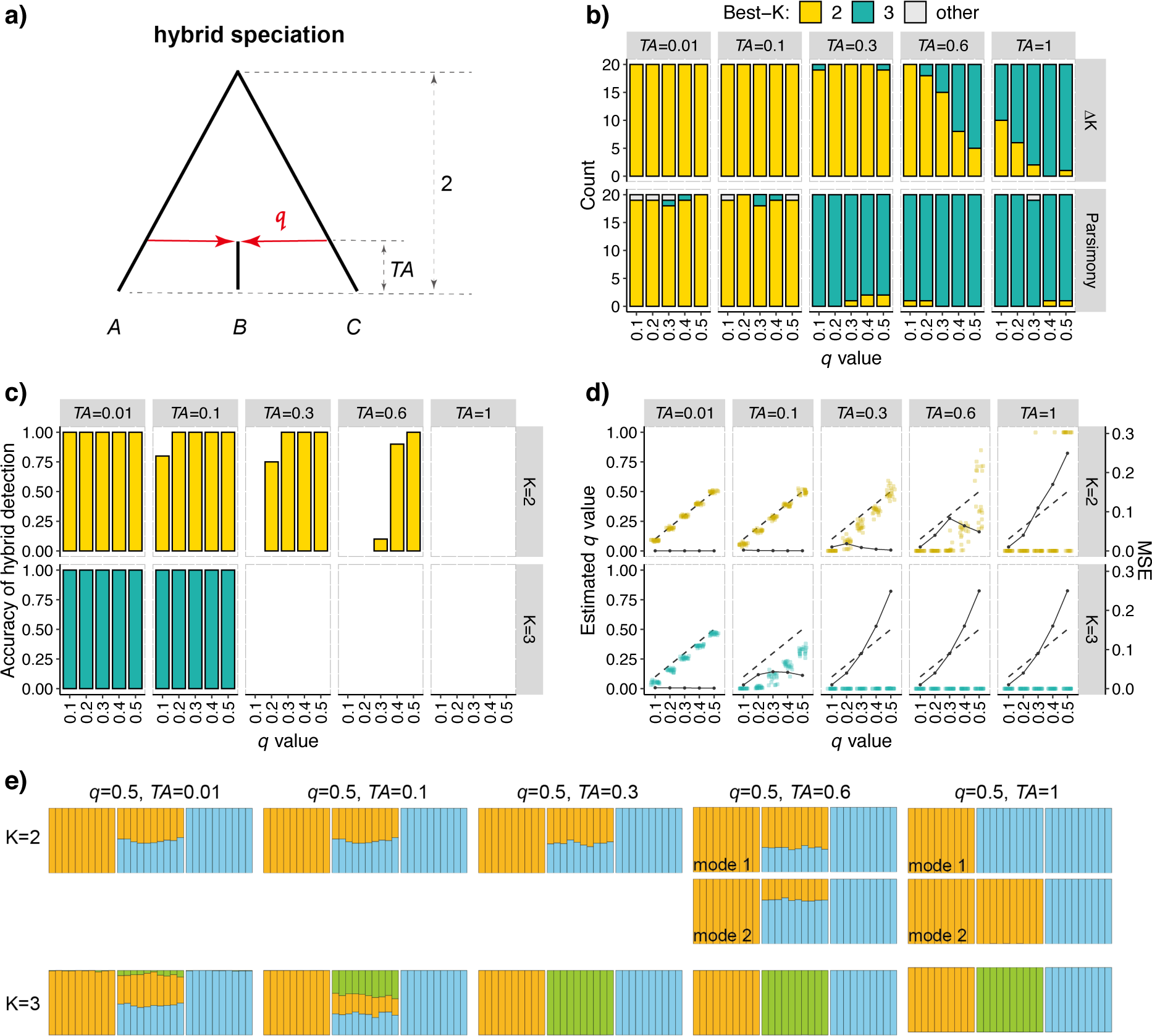
Results for the scenarios of hybrid speciation. a) Hybrid speciation scenarios with corresponding parameter settings. *TA* represents the time of the admixture event (in coalescent units), and *q* represents the proportion of genetic material in hybrids inherited from parent *C*. b) Best-*K* inference using Δ*K* and parsimony methods. The colored bars represent the numbers of inferences for best-*K* = 2 and best-*K* = 3 among 20 replicates. The strips located at the top and right of each plot represent *TA* and the methods employed, respectively. The x-axis indicates the values of *q*. c-d) Plots of the results for hybrid detection and estimation of admixture proportion (*q*-value) when assuming *K* = 2 and *K* = 3. The strips located at the top and right of each plot represent *TA* and the assumed number of clusters K, respectively. c) Accuracy of hybrid detection. The y-axis represents the proportion of times that hybrids were successfully identified. d) Estimation of admixture proportion *q*-value. Colored points represent the average estimates of *q* for hybrid individuals within each dataset, and are horizontally jittered to avoid clutter. The dashed line shows the true values, while the solid line with points represents the MSE values. e) Inferred STRUCTURE clustering plots for *K* = 2 and *K* = 3 when *q* = 0.5. At the top of each plot, the specific introgression scenario and the parameter settings (*q* and *TA*) are indicated. Each plot illustrates the modes of clustering for individuals from species *A*, *B*, and *C* (from left to right). For additional clustering plots for other parameters, refer to Figure S1.

**FIGURE 2.**
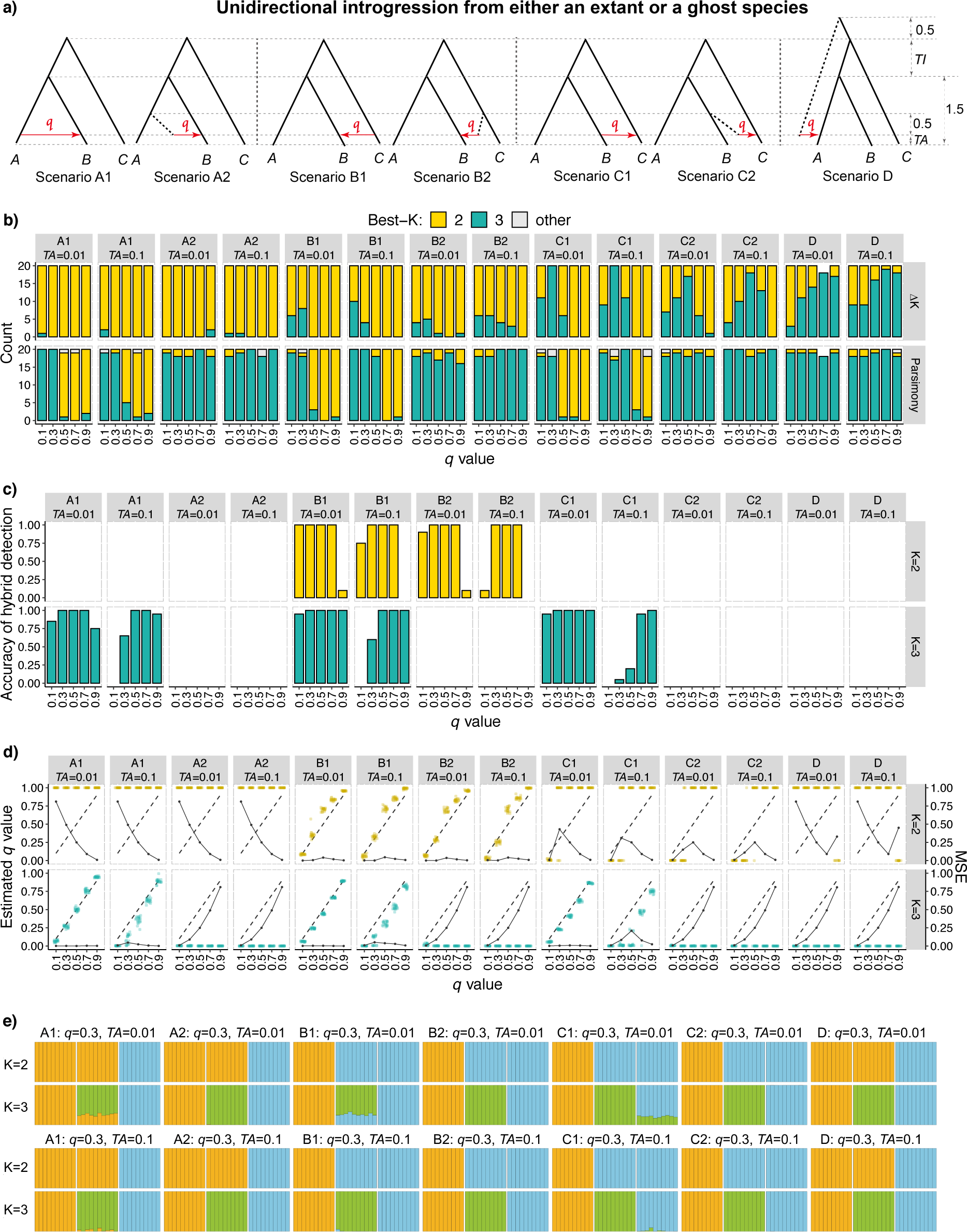
Results for scenarios involving unidirectional horizontal introgression or ghost introgression with *TI* = 0.5. a) Simulated introgression scenarios and parameter settings. A1—Introgression between sister species (𝐴 → 𝐵); A2—Ingroup ghost introgression from an *A*-derived ghost to *B*; B1—Inflow introgression (𝐶 → 𝐵); B2—Ingroup ghost introgression from a *C*-derived ghost to *B*; C1—Outflow introgression (𝐵 → C); C2—Ingroup ghost introgression from a *B*-derived ghost to *C*; D—Outgroup ghost introgression. *TA* represents the time of the admixture event (in coalescent units) and *TI* represents the time interval between two speciation events within the species tree (in coalescent units). *q* signifies the proportion of genetic material in hybrids inherited from the donor species. b) Best-*K* inference using Δ*K* and parsimony methods. Colored bars represent the numbers of inferences for best-*K* = 2 and best-*K* = 3 among 20 replicates. The strip at the top of each plot indicates the simulated scenarios and the parameter *TA*, while the strip on the right of each plot represents the methods employed. The x-axis indicates the values of *q*. c-d) Plots of the results for hybrid detection and estimation of admixture proportion (*q*-value) assuming *K* = 2 and *K* = 3. The strips on the right of each plot represent the chosen number of clusters *K*. The x-axis indicates the values of *q*. c) Accuracy of hybrid detection. The y-axis represents the proportion of times that hybrid individuals were successfully identified. d) Estimation of admixture proportion *q*-value. Colored points represent the average estimates of *q* for hybrid individuals within each dataset, and are horizontally jittered to avoid clutter. The dashed line shows the true values, while the solid line with points represents the MSE values. e) Inferred STRUCTURE clustering plots for *K* = 2 and *K* = 3 when *q* = 0.3. At the top of each plot, the specific introgression scenario and the parameter settings (*q* and *TA*) are indicated. Each plot illustrates the modes of clustering for individuals from species *A*, *B*, and *C* (from left to right). For clustering plots for other parameter combinations, refer to Figures S4-S10.

**FIGURE 3.**
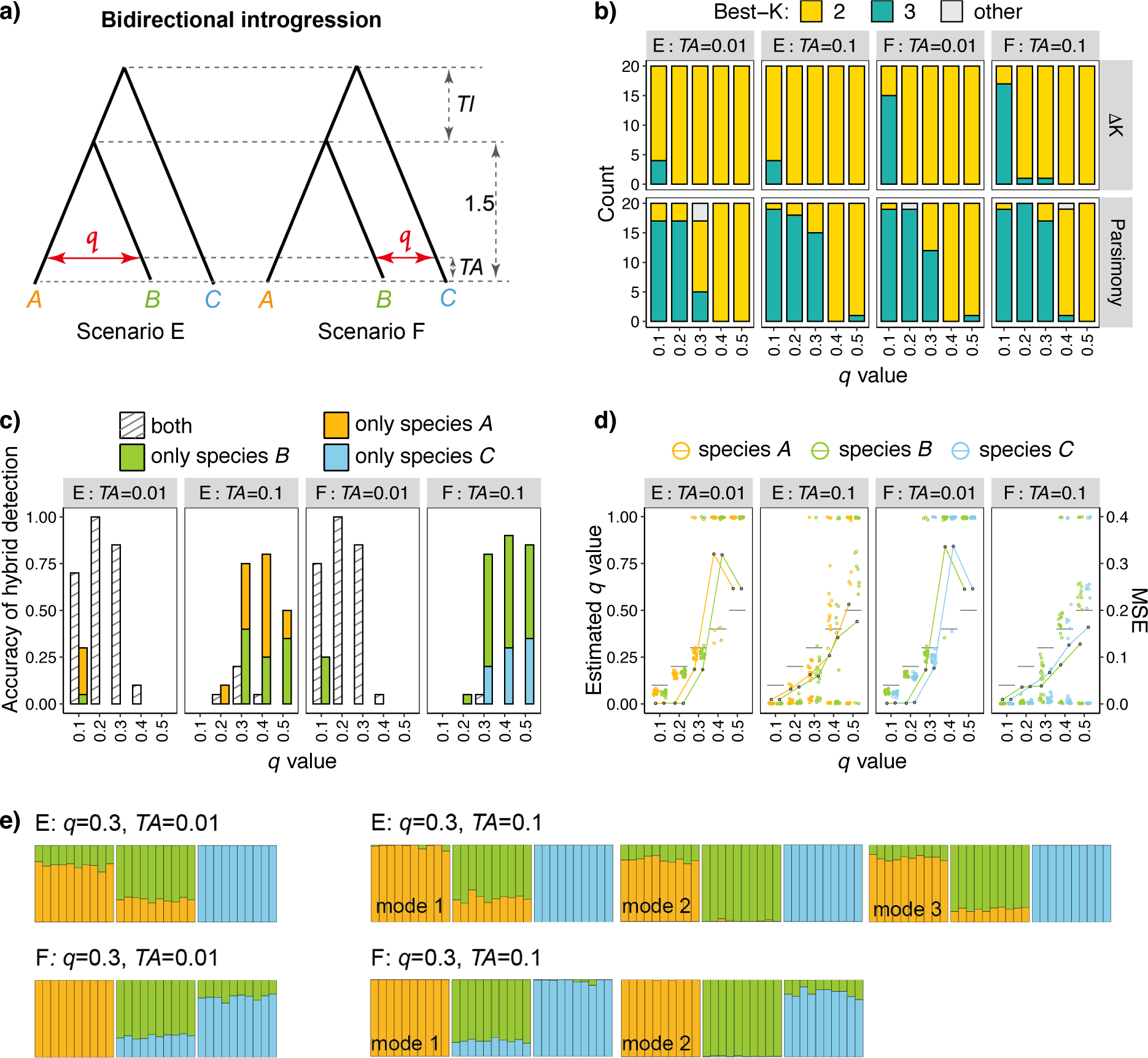
Results for bidirectional introgression scenarios with *TI* = 0.5. a) Introgression scenarios with corresponding parameter settings. Introgression levels between species in different directions are assumed to be symmetrical and are labeled by *q*. *TI* represents the time interval between two speciation events (in coalescent units) within the species tree. b) Best-*K* inference using Δ*K* and parsimony methods. Colored bars represent the numbers of inferences for best-*K* = 2 and best-*K* = 3 among 20 replicates. The strip at the top of each plot indicates the simulated scenarios and the parameter *TA*, while the strip on the right of each plot represents the methods employed. The x-axis indicates the values of *q*. c-d) Plots of results for hybrid detection and admixture proportion estimation assuming *K* = 3. The results for *K* = 2 are omitted for brevity, as STRUCTURE with *K* = 2 failed to identify any hybrids. Results for different hybrid species are labeled in different colors. c) Accuracy of hybrid detection. The y-axis shows the proportion of times that hybrid lineages are successfully identified. The legend categorizes detection results into four distinct types: ’both’—concurrent identification of two hybrid lineages; ’only species *A*’, ’only species *B*’ and ’only species *C*’—a single hybrid lineage is identified, corresponding to either species *A*, *B*, or *C*, respectively. d) Estimation of admixture proportion *q*-values. Colored points represent the average estimates of *q*-values for hybrid individuals from each hybrid lineage across the datasets, horizontally jittered to avoid clutter. Black lines indicate the true *q*-values, and colored lines with black points represent MSE values. e) Inferred STRUCTURE clustering plots for *K* = 3 when *q* = 0.3. At the top of each plot set, the specific introgression scenario and the parameter settings (*q* and *TA*) are indicated. Each set of plots illustrates clustering modes for individuals from species *A*, *B*, and *C* (from left to right). For clustering plots for other parameter combinations, refer to Figures S13-S14.

## Materials And Methods

### Simulation Settings

We generated simulation data for various scenarios under the model of multispecies coalescent with introgression (Yu et al. 2014; MSci), which assumes episodic gene flow occurring at a fixed time point in the past. We kept the effective population size constant at N = 10,000 for all extant and ancestral species. The times of evolutionary events were measured in coalescent units, where one coalescent unit is equivalent to 2N or 20,000 generations. For scenarios of hybrid speciation (Fig. 1a), we set the divergence time between the two parental species at 2 but varied the time of the admixture event (*TA*) to be 0.01, 0.1, 0.3, 0.6, and 1. The contribution proportion in the hybrid lineage *B* from species *C* (*q*) was set to be 0 (i.e., no introgression), 0.1, 0.2, 0.3, 0.4, and 0.5.

For all scenarios of introgression occurring within a fixed species tree *AB|C* (Figs. 2a and 3a), we kept the divergence time between sister species at a fixed value of 1.5. The length of the internal branch, denoted as *TI*, was set to be either 0.5 or 1.5. We investigated two distinct admixture times *TA*, with values of 0.01 and 0.1, and set a range of proportions of genetic contribution (*q*) from the donor species, specifically 0.1, 0.3, 0.5, 0.7, and 0.9. For each case of unidirectional horizontal introgression, we adjusted the reticulation edge length to 0.5 to simulate the corresponding ingroup ghost introgression. In scenarios of outgroup ghost introgression (Scenario D in Fig. 2a), the distance of the outgroup ghost to the most recent common ancestor of the three focal species was set to 0.5.

Simulations for scenarios of hybrid speciation and unidirectional introgression were run using the program *ms* (Hudson 2002) with the option -s 1 to generate single nucleotide polymorphism (SNP) loci. For scenarios with bidirectional introgression, we utilized the *simulate* option of BPP version 4.7 (Rannala and Yang 2017; Flouri et al. 2023) to generate sequences under the JC69 model (Jukes and Cantor 1969). Subsequently, we utilized a custom Perl script to process the sequence data, selectively retaining a single SNP from each locus to create independent SNP datasets for STRUCTURE analysis. In each case, each species comprised 10 diploid samples. 20 replicate datasets were simulated, each comprising 1000 independent SNPs unless otherwise specified.

### STRUCTURE Analyses and Evaluation

STRUCTURE utilizes an admixture model to identify hybrid individuals (Pritchard et al. 2000). Specifically, from a postulated number of clusters *K* (interpreted as source genetic pools), this model characterizes the genetic composition of individuals by estimating the genetic contribution proportions, *q*, from *K* clusters. In practice, *q*-values are commonly used for classifying individuals as pure or admixed by applying a threshold (e.g., 0.9 or 0.95). If the *q*-value from a cluster exceeds the threshold, the individual is classified as a pure member of that cluster; otherwise, it is categorized as admixed.

For each dataset, we ran STRUCTURE with both the admixture and correlated allele frequencies models, as recommended in the STRUCTURE manual. The analyses were carried out for *K* values ranging from 1 to 4, with each *K* being subjected to 5 independent runs. Following Kong and Kubatko (2021), who analyzed larger datasets than ours, we configured the parameters for the length of the burn-in period and the number of MCMC iterations to 50,000 and 100,000, respectively. Besides, we assessed two best-*K* inference methods: the widely-used Δ*K* method (Evanno et al. 2005) and a recently proposed parsimony estimator, *PI* (Wang 2019). Both methods were implemented using the KFinder software tool (Wang 2019). We paid special attention to the results of STRUCTURE for *K* = 2 and *K* = 3, as these were the two *K* values most frequently suggested by the best-*K* inference methods in our results, spanning all scenarios considered. For each data replicate, we selected the run with the highest posterior probability to represent the most accurate estimation.

Our evaluation centers on two key aspects of STRUCTURE’s performance: the identification of hybrid lineages and the estimation of contribution proportions from donor species. We applied a relatively stricter *q*-value threshold of 0.95 for hybridization detection in consideration of the minimum introgression level set at 0.1 in our simulation settings. We treated hybrid lineages as the unit of assessment, calculating the average admixture proportions (*q*) for all individuals within a lineage. The accuracy of hybrid detection was measured as the proportion of times that true hybrid lineages were classified as admixed across replicates. The precision of estimated contribution proportions from donors was assessed by calculating the mean squared error (MSE) across replicates. The MSE is calculated by the formula: 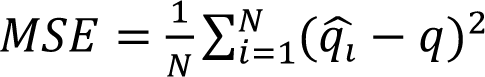, where *N* represents the number of replicates (i.e., 20), *q* is the true value of the contribution proportions from donors as specified in the simulation, and 𝑞(_#_ is the estimate value of *q* for the *i*th replicate.

## Results

### Hybrid Speciation

The results from various cases of hybrid speciation, which involve different combinations of admixture proportions (*q*) and admixture times (*TA*), are summarized in Figure 1. When it comes to inferring the optimal number of clusters (best-*K*), the Δ*K* method indicated the best-*K* of 2 when the admixture event was relatively recent (i.e., *TA* ⩽ 0.3). However, when *TA* ⩾ 0.6, the results were less consistent across multiple data replicates; some replicates suggested a best-*K* of 2, while others suggested 3 (as seen in Fig. 1b). In contrast, the parsimony method was more sensitive to admixture timing and showed greater stability, suggesting that the best-*K* is 2 for *TA* ⩽ 0.1, and 3 for higher values of *TA*.

Furthermore, we delved into the behaviors of STRUCTURE when specifying *K* = 2 and *K* = 3, respectively. When setting *K* = 2, for recent admixture events at *TA* ⩽ 0.1, STRUCTURE effectively identified both hybrid lineage *B* and its parents *A* and *C*, as well as precisely estimated the admixture proportions as reflected by their small MSE values (Figs. 1c-e and S1). However, as *TA* increased, the estimated admixture proportions became progressively more deviant from the true value, that is, a kind of ‘the Matthew effect’ (Fig. 1d): for the minor parent with *q* < 0.5, its admixture contribution was underestimated, whereas for the major parent with *q* > 0.5, it was overestimated. Consequently, admixture events with asymmetrical parental contributions are more difficult to detect, a challenge also highlighted in the simulation study of Kong and Kubatko (2021). Interestingly, when the parental contribution was symmetrical (*q* = 0.5), the estimated *q*-values began to exhibit a bimodal distribution as the time of admixture *TA* increased to above 0.6 (Fig. 1e). If the hybrid lineage was formed as moderately ancient as *TA* = 1, admixture signals may be lost altogether, with the hybrids erroneously being grouped with one of their parents (Figs. 1e). In practical terms, this means that symmetrical parental contribution will often be revealed as extremely asymmetrical in STRUCTURE, especially when the hybrid lineage was formed in a relatively long past.

If setting *K* = 3, STRUCTURE successfully identified the hybrids only for very recent admixture events with *TA* ⩽ 0.1, but the donor lineages were not inferred correctly. To be more specific, there are three ancestry components for the hybrid lineage, including the two actual parents, as well as a shadowy gene pool that has no purebred members being recognized in the sample. For moderately ancient admixture events with *TA* ⩾ 0.3, the admixed individuals, as expected, were identified as a third independent cluster (Fig. 1e and S1).

Figures 2 and S3 illustrate the outcomes of unidirectional introgression under various conditions. These conditions include the presence or absence of ghost lineages and variations in key parameters: admixture times (*TA*), time intervals between speciation events (*TI*), and admixture proportions (*q*). In terms of best-*K* inference, the Δ*K* method consistently leaned towards endorsing *K* = 2 across all introgression scenarios except for outgroup ghost introgression (Figs. 2b and S3a), whereas the parsimony method tended to favor *K* = 3 as *TA* increased and/or as *q* decreased in scenarios of horizontal introgression (Scenarios A1, B1, and C1). However, when introgression originated from a ghost lineage as depicted in scenarios A2, B2, C2, and D, the parsimony method consistently inferred the best-*K* to be 3 across all combinations of parameters. In contrast, the influence of ghost lineages on the Δ*K* method was less pronounced, suggesting best-*K* = 2 for most of *q* ⩾ 0.5 in scenarios (A2, B2 and C2) of ingroup ghost introgression where the extinct or unsampled donor (ghost) diverged within the timeframe since the most recent common ancestor of the sampled species.

Then, we take a closer look at the behavior of STRUCTURE with *K* = 2 or *K* = 3, respectively. In the absence of introgression within the species tree *AB|C* (i.e., *q* = 0), STRUCTURE yielded perfect results as expected (Fig. S2). Specifically, with *K* = 2, individuals from sister species *A* and *B* were first grouped into one cluster, while another cluster comprised individuals from the early-diverged species *C*. When *K* was increased to 3, STRUCTURE accurately assigned individuals to their respective species.

Moving on to horizontal or non-horizontal (ghost) introgression scenarios, we observed the following outcomes. Firstly, when *K* was set to 2, STRUCTURE, as expected, grouped individuals of sister species into a single cluster in scenarios of sister introgression (i.e., Scenarios A1 and A2) (Figs. S4-S5). Surprisingly, in cases of inflow introgressions (i.e., Scenario B1 and B2), STRUCTURE with *K* = 2 could sometimes identify the hybrid lineage *B*, with the success rate being much higher under moderate ILS (*TI* = 1.5; Fig. S3) than under prevalent ILS (*TI* = 0.5; Fig. 2c). The successful identification of hybrid lineages in cases of horizontal inflow introgression was also observed in the study of Kong and Kubatko (2021). However, in these cases, the sister species (*A*) of the hybrid lineage (*B*) was incorrectly diagnosed as a parental lineage (Figs. S6-S7), which, in turn, led to an inflated estimate of introgression proportions from the actual donor species *C* (Figs. 2d and S3c). In non-horizontal inflow introgression scenarios, where gene flow occurred from a *C*-derived ghost to species *B* (i.e., Scenario B2), STRUCTURE further erroneously inferred the participation of species *C* in gene flow as a donor, incorrectly implying a horizontal reticulation event (Figs. S6-S7). In contrast, for outflow introgression with an opposite gene flow direction (Scenarios C1 and C2) and outgroup ghost introgression (Scenario D), STRUCTURE with *K* = 2 failed to detect any hybrids. Instead, the trend shifted from initially grouping sister species *A* and *B* together to grouping non-sister species *B* and *C* together as the level of introgression increased (Figs. S8-S10).

When the correct number of clusters (*K* = 3) was specified to represent the gene pools of the three sampled species prior to gene flow, STRUCTURE efficiently identified both hybrid lineages and the donors of introgression, as well as providing reasonably accurate estimates of introgression levels (*q*) for all scenarios of horizontal introgression (i.e., Scenario A1, B1, and C1) when admixture was very recent (*TA* = 0.01) (Figs. 2c, S3b). However, STRUCTURE is much sensitive to post-admixture genetic drift, with an obvious decrease in the accuracy of hybrid detection even for moderately recent admixture (*TA* = 0.1), particularly when introgression intensity was not too high (*q* ⩽ 0.5) (Figs. 2c, S3b). More specifically, when *TA* = 0.1, the contribution proportions (*q*) from donors were consistently underestimated across all horizontal introgression scenarios, exhibiting three distinct underestimation patterns depending on the gene flow level (Figs. 2d and S3c). For weak introgression events (*q* ⩽ 0.1), the admixture proportions *q* were underestimated to near zero, leading to a failure in detecting hybridization signals. In contrast, for intense introgression events (*q* ⩾ 0.7), the contribution proportions from donors were underestimated but remained above 0.5, which were sufficient to identify hybrid individuals. Intriguingly, in cases of intermediate levels of introgression (*q* = 0.3 or 0.5), two modes were observed among replicates: some replicates showed the estimated *q* values approaching 0, thereby failing to detect hybridization, while others exhibited a relative minor underestimation, allowing for the detection of hybrids. Besides, the extent of sensitivity to the timing of admixture is linked to the degree of ILS and the direction of introgression, with a notable decrease in hybrid detection accuracy in cases with substantial ILS (*TI* = 0.5) or outflow introgression as the admixture time (*TA*) increases (Figs. 2c, S3b).

Most surprisingly, when investigating introgression originating from a ghost lineage— regardless of whether the ghost lineage diverged prior to or after the root of the species tree (i.e., ingroup versus outgroup)—STRUCTURE with *K* = 3 consistently failed to detect hybridization. It incorrectly classified hybrid individuals as purebred across all tested parameters of *TI*, *TA*, and *q* (Figs. 2 and S3).

Bidirectional introgression is widely recognized as a common form of genetic interchange between species (Yang and Flouri 2022). The two species involved in gene flow serve as both donors and recipients to each other. Thus, bidirectional introgression is horizontal introgression by definition. In this section, we explore how well STRUCTURE performs in such introgression scenarios. Our analysis was limited to cases with symmetrical introgression levels in both directions. In consideration of the unidentifiability between the parameter *q* and 1-*q* in this model (Fig. S11; Yang and Flouri 2022), we set the contribution proportion *q* from the donor species to be 0.1, 0.2, 0.3, 0.4, and 0.5. The results are presented in Figures 3 and S12.

For the best-*K* inference, the Δ*K* method generally endorsed *K* = 2 across the tested cases, except for situations involving introgression between non-sister species at *q* = 0.1 (Figs. 3b and S12b). By contrast, the parsimony method inferred a best-*K* of 3 for most moderate introgression cases of *q* ⩽ 0.3.

When setting *K* = 2, individuals from the two species engaged in introgression were generally grouped into a single cluster (Figs. S13-S14). An exception was observed in the scenario of introgression between non-sister species with a moderate level of ILS at *TI* = 1.5 and a low level of introgression at *q* = 0.1 (Fig. S12b), in which only one species was successfully identified as admixed. But STRUCTURE erroneously attributed the genetic donors of the admixed species *B* to species *A* and *C*, despite species *A* not being involved in the gene flow event and species *C* being itself an admixed lineage (Fig. S14).

When we set the number of clusters as *K* = 3, in cases of very recent introgression at *TA* = 0.01, STRUCTURE could identify two hybrid species simultaneously with an accuracy exceeding 60% when *q* ⩽ 0.3 (Figs. 3c and S12b), regardless of whether gene flow took place between sister or non-sister species. In these cases, admixture proportion *q* for the detected hybrids tended to be slightly underestimated (Figs. 3d and S12c). When *q* ⩾ 0.4, all individuals from the two species engaged in the introgression were assigned to the same cluster, similar to what was already observed with *K* = 2 (Figs. S13-S14).

In cases where introgression occurred in comparatively deeper time at *TA* = 0.1, STRUCTURE mostly failed to reveal any hybrids when *q* ⩽ 0.2. In scenarios of sister introgression at *q* ⩾ 0.3, signals of hybridization were detected, yet they manifested in various admixture patterns (i.e., multiple clustering modes). Occasionally, both hybrid lineages—species *A* and *B*—were identified concurrently, indicating bidirectional introgression; however, more frequently, only one hybrid lineage was detected, implying unidirectional introgression from *A* to *B* or the other way around (Figs. 3c, 3e and S12b). When introgression occurred between non-sister species with *q* ⩾ 0.3, STRUCTURE preferred to solely identify species *B* as an admixed lineage, especially under conditions of moderate ILS with *TI* = 1.5 (Figs. 3c, 3e and S12), which, in turn, incorrectly suggested a pattern of unidirectional inflow introgression. Moreover, when *TA* = 0.1, the accuracy and precision of the estimated *q* were relatively poor, with *q* values being significantly underestimated at *q* = 0.3 and overestimated at *q* = 0.5 (Figs. 3d and S12c).

### Outgroup Ghost Introgression under Continuous Gene Flow

We have used the MSci model of gene flow so far,, which assumes that cross-species gene flow occurs at a fixed time in the past. Another class of gene flow model has also been developed, known as the isolation-with-migration model (IM; Hey and Nielsen 2004), which assumes that two species exchange migrants at a certain rate over an extended time period. Here, we extended our analysis from episodic to continuous gene flow to assess if our results obtained with STRUCTURE for the detection of ghost introgression are sufficiently general. Specifically, we examined scenarios of outgroup ghost introgression, with continuous migration spanning from the divergence of sister species to the present (Fig. S15a).

We explored a range of migration rates *M* = 0.4, 0.8, 1.6, 3.2, and 6.4, with outcomes presented in Figure S15. When setting *K* = 2, STRUCTURE grouped individuals of the sister species *A* and *B* into one cluster for small migration rates of *M* ⩽ 1.6, whereas it grouped non-sister species *B* and *C* together for the larger *M* values of ⩾ 3.2. When *K* = 3, individuals from all three species were consistently segregated into separate clusters, irrespective of *M* values (Fig. S15b). In brief, when the gene flow stems from ghost lineages through continuous migration, STRUCTURE still fails to detect hybridization signals across all tested parameter combinations and irrespective of the chosen *K* values. These findings reconfirm the above results concerning STRUCTURE’s inability to detect hybridization when donor lineages are not included in the sample for analysis.

### Effect of Number of SNPs

In this section, we investigated the effect of SNP numbers on STRUCTURE’s performance, with a specific focus on addressing two questions: (i) Can increasing SNP numbers enhance STRUCTURE’s ability to detect deep reticulation or ghost introgression? (ii) Can increasing SNP numbers overcome the observed problem of multiple clustering modes in our results? To explore these questions, we focused on cases of horizontal or non-horizontal (ghost) introgression between sister species *A* and *B* with *TI* = 0.5 and *q* = 0.3. We chose the admixture times *TA* as either 0.01 or 0.1, and we analyzed three different datasets consisting of 250, 1000, 4000, and 16000 SNPs. The results of this analysis, conducted using STRUCTURE with *K* = 3, are presented in detail in Figure 4.

**FIGURE 4.**
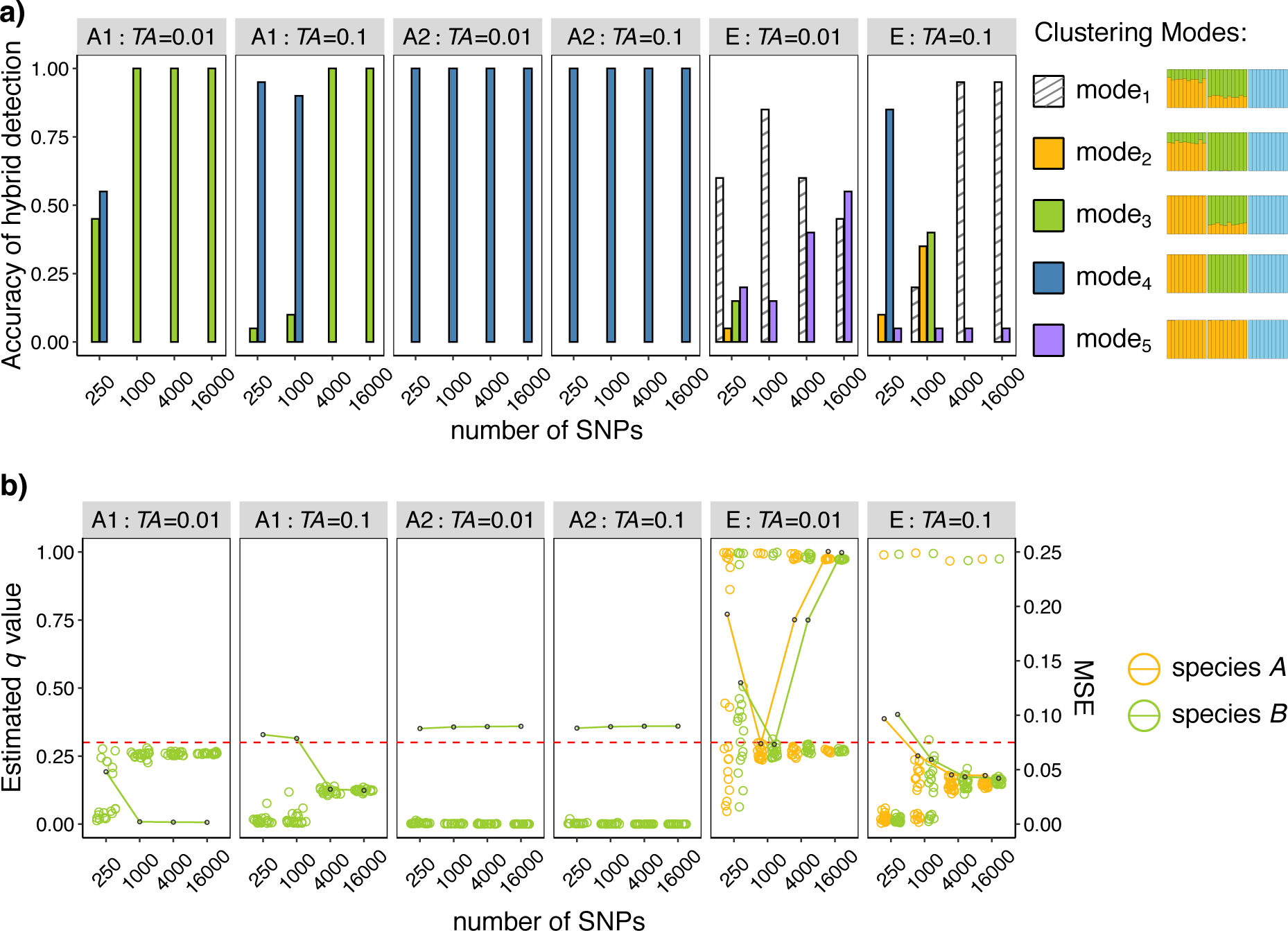
The effect of the number of SNPs on STRUCTURE’s performance. The scenarios considered include unidirectional horizontal, non-horizontal (ghost), and bidirectional introgression between sister species (i.e., Scenarios A1 and A2 in Fig. 2a, and Scenario E in Fig. 3a). The analysis was limited to cases with the parameters *q* = 0.1 and *TI* = 0.5, conducted using STRUCTURE with *K* = 3. The strip at the top of each plot indicates simulated scenarios and the time of introgression *TA* (in coalescent units). The x-axis shows the number of SNPs. a) Inferred different clustering modes across replicates. b) Estimation of admixture proportion *q*-values. Colored points represent the average estimates of *q*-values for hybrid individuals from each hybrid lineage across 20 replicate datasets. The dashed red lines indicate the true *q*-values, and colored lines with black points represent MSE values.

Our results indicated that increasing the number of SNPs enhanced the ability to detect historical horizontal introgression. However, as introgression events became older, the number of SNPs required for reliable detection escalated significantly (Fig. 4). For example, in instances of very recent unidirectional horizontal introgression (i.e., Scenario A1) at *TA* = 0.01, a dataset of 1,000 SNPs sufficed to identify introgression. In contrast, moderately old admixture events at *TA* = 0.1 necessitated as many as 4,000 SNPs for detecting admixture signals, yet the estimated *q*-values were substantially underestimated, registering at around 0.12, well below the true 0.3 (Fig. 4b). Considering the marked trend of *q*-value underestimation with increasing the time of admixture, it is reasonable to expect that for deeper reticulation events, even a substantial SNP dataset may be ineffective if the best-fitting *q*-values fall below the designated threshold (e.g., 0.05). When introgression was derived from a ghost lineage (i.e., Scenario A2), STRUCTURE invariably yielded estimates of negligible *q*-values, independent of the size of the used datasets (Fig. 4).

The issue of multiple clustering modes cannot necessarily be resolved by increasing the number of SNPs. For instance, in scenarios of very recent bidirectional introgression (i.e., Scenario E) at *TA* = 0.01, both the correct mode (mode_1_), where both sister species *A* and *B* show admixed, and the false mode (mode_5_), where both sister species appear as pure and grouped in the same cluster, persistently occurred regardless of the specific SNP dataset used (Fig. 4a). Notably, when the number of SNPs was increased from 1000 to 16000, the proportion of the false mode noticeably increased to 55%, comparable to that of the correct mode (45%). The counterintuitive performance deterioration upon adding more SNPs suggests that in some cases, “less is more,” meaning that a smaller number of SNPs may actually lead to better results.

## Discussion

Population clustering methods, which many researchers had long cherished for detecting hybridization on deep evolutionary scales (Blair and Ané 2020; Stull et al. 2023), have yet to be thoroughly evaluated for their effectiveness in such contexts. Our investigation addresses this knowledge gap by systematically evaluating the performance of STRUCTURE under a variety of hybridization scenarios on deep time-scales, including hybrid speciation and horizontal as well as non-horizontal (ghost) introgression. We place particular emphasis on the effects of admixture timing and ghost lineages—common confounding factors encountered in longer time periods—on hybridization detection. Our findings reveal that STRUCTURE struggles to detect admixture signals when hybridization occurs in deep time or stems from ghost lineages. Among numerous model-based clustering algorithms that have been developed so far (reviewed in Novembre 2016; Lawson et al. 2018), STRUCTURE is the first and most popular. Notably, STRUCTURE has been shown to be more powerful than ADMIXTURE (Alexander et al. 2009)—another popular likelihood-based approach with improved computational speed—in terms of hybridization detection (Kong and Kubatko 2021). Thus, it is reasonable to presume that the same caveats would generally apply if any population genetic clustering approach is used to detect hybridization across phylogenetic time-scales.

### Deep Reticulation: Challenges in Detection Using Population Clustering Methods

For very recent horizontal hybridization events occurring between sister or non-sister species, STRUCTURE proves to be highly effective in accurately identifying hybrid individuals and pinpointing their donor lineages. It also provides reasonably accurate estimates of contribution proportions from donors (*q*), albeit with a slight tendency to underestimate in most cases. Additionally, STRUCTURE can discern recent bidirectional introgression events when introgression is not very extensive (i.e., *q* ⩽ 0.3 in Figs. 3 and S12). This ability to identify the direction of recent introgression, even between sister species, stands out as a significant advantage of population clustering methods. This contrasts with many phylogenetic approaches, which typically cannot ascertain the direction of introgression and may even fail to detect hybridization between sister lineages (Hibbins and Hahn 2022). Such an advantage of population clustering methods derives from their utilization of population-level data from multiple samples per species, as opposed to phylogenetic methods that often rely on a single sample per species (Hibbins and Hahn 2022). However, it is crucial to recognize that when the dataset is limited to a small number of markers, such as a few hundred SNPs, there can be a considerable underestimation of admixture levels (*q*), severely compromising STRUCTURE’s hybrid detectability (Fig. 4). The adverse impact of using a small number of markers on population clustering methods has also been highlighted by Toyama et al. (2020). This underscores the necessity of employing large-scale datasets, including whole-genome data whenever possible, to enhance the detectability of population clustering methods.

Identifying ancient reticulation events, however, poses a significant challenge for population clustering methods. Our results indicate that STRUCTURE’s ability to detect hybridization is highly sensitive to the time elapsed since admixture (Figs. 1-3). This sensitivity arises due to post-admixture genetic drift altering allelic frequencies in both hybrid and donor species, thereby eroding the hybridization signal (Toyama et al. 2020). For instance, in scenarios of hybrid speciation with symmetrical parental contributions (*q* = 0.5)—a common expectation if reproductive isolation begins immediately in F1 hybrids (Hibbins and Hahn 2019)—when the hybrid lineage originates in the moderately distant past (e.g., with the admixture time *TA* of 0.6 coalescent units, compared to the speciation time of the parental lineages at 2 coalescent units), the estimated *q*-values tend to exhibit a bimodal distribution, clustering both above and below the true value of 0.5 (Fig. 1e). And as the admixture event becomes more ancient, the estimated *q*-values increasingly gravitate towards the extreme values of 0 and 1, indicating a growing asymmetry in parental contributions and eventually resulting in a complete loss of admixture signals (e.g., at *TA* = 1 in Fig 1e). Practically speaking, symmetrical parental contributions to the hybrid lineage are often misleadingly revealed as asymmetrical using population clustering methods, particularly when admixture happened in a long past (say, tens of thousands of generations ago).

Ancient cross-species introgression is even more challenging to detect using population clustering methods. Compared with hybrid speciation, post-admixture genetic drift plays a more significant role in distorting the estimation of admixture proportions in cases of introgression, leading to a greater likelihood of failure in identifying hybridization when the admixture event is not very recent. For instance, in cases where two species that have diverged for 2 coalescent units hybridized at *TA* = 0.1, STRUCTURE performs well for hybrid speciation scenarios (Fig. 1c) but struggles to identify hybrids in unidirectional introgression scenarios when the admixture level is not high (i.e., *q* ⩽ 0.3 in Fig. 2c). Additionally, even in cases with prevalent hybridization (*q* ⩾ 0.3 in Fig. 3c), STRUCTURE tends to misidentify relatively ancient bidirectional introgression as unidirectional. Given that hybridization is more likely to occur in early stages of speciation process, population clustering methods would inevitably lead to a substantial underestimation of the incidence of historical introgression.

Furthermore, the direction of introgression plays an important role in detecting historical introgression between non-sister species. For example, in cases of unidirectional introgression that is not very recent (e.g., *TA* = 0.1), STRUCTURE exhibits higher accuracy in detecting inflow introgression (from the outgroup species to an ingroup species) than outflow introgression (Figs. 2c and S3b). More interestingly, in scenarios involving bidirectional introgression between non-sister species with *TA* = 0.1, STRUCTURE tends to predominantly detect the inflow introgression while largely missing the outflow introgression (Fig. 3c). These findings consistently demonstrate that inflow introgression is more readily detectable compared to outflow introgression. Importantly, such a pattern is not confined to population clustering methods; it also applies to other popular approaches for detecting introgression (Zheng and Janke 2018; Thawornwattana et al. 2023; Pang and Zhang 2024), such as D-statistics (Green et al. 2010), HyDe (Blischak et al. 2018), and BPP (Flouri et al. 2020). The consistency across these various approaches implies that inflow introgression leaves behind a richer set of genetic signals than outflow introgression, making it more amenable to detection.

### Ghost Lineage: An Important Factor Further Complicating Hybridization Detection through Admixture Analysis

Ghost introgression, a widespread phenomenon in nature, has become a pivotal consideration in evolutionary research in recent years (Ottenburghs 2020; Tricou et al. 2022a, 2022b; Pang and Zhang 2023, 2024). Several studies have underscored the significance of considering ghost lineages when investigating hybridization (Lawson et al. 2018; Garcia-Erill and Albrechtsen 2020; Tricou et al. 2022a, 2022b; Pang and Zhang 2024). Notably, Lawson et al. (2018) found that the extinction of parental lineages could result in misattributed sources of gene flow in admixture analyses. This raises concerns about the potential broader impacts of ghost lineages on population genetic clustering techniques (Tricou et al. 2022a). By examining a wider range of ghost introgression scenarios, our study provides the first comprehensive analysis of the potential effects of ghost lineages on hybridization detection through population clustering methods.

Our results demonstrate that population genetic clustering methods have inherent difficulties in detecting ghost introgression. If introgression originates from a ghost lineage— regardless of the position of the ghost lineage (ingroup versus outgroup), the timing of admixture (recent versus ancient), and the mode of gene flow (continuous versus episodic)— STRUCTURE almost always fails to identify hybrid individuals (Figs. 2 and S3). Importantly, this failure appears unsolvable by increasing the sample size of SNPs (Fig. 4). These results show that the absence of introgression donors in the data may make admixture signals undetectable with population clustering methods.

However, there is a noteworthy caveat to consider: in the scenario where introgression occurs from a *C*-derived ghost to one of the two sister species (Scenario B2 in Fig. 2), STRUCTURE with *K* = 2 does successfully identify hybrid individuals under moderate ILS (*TI* = 1.5; Fig. S3), but it misidentifies the donor as the sampled lineages. This result could be misinterpreted as horizontal gene flow. Similar misidentification of donor species when hybridization involves ghost lineages has also been reported in previous population-level studies by Lawson et al. (2018) and Garcia-Erill and Albrechtsen (2020).

Overall, within the context of ghost introgression, population clustering methods either fail to detect the admixture signals or misidentify them as horizontal gene flow. This partly explains why ghost introgression has escaped the attention of evolutionary biologists until recently. As ghost introgression receives increasing attention (Lawson et al. 2018; Garcia-Erill and Albrechtsen 2020; Tricou et al. 2022a, 2022b; Pang and Zhang 2023, 2024), researchers are intensifying their efforts to investigate its potential effects and explore new methods to detect and quantify it, which would prompt a reexamination of past overlooked ghost introgression events.

### Recommendations for Empirical Studies

*On the selection of the best-K (number of clusters)*—Across various scenarios involving three species considered in this study, the parsimony method (Wang 2019) tends to infer the optimal number of clusters as *K* = 3, while the Δ*K* method (Evanno et al. 2005) often shows a preference for *K* = 2. This aligns with the earlier finding from Janes et al. (2017), which showed that *K* = 2 was notably overrepresented in empirical studies using the Δ*K* method. For horizontal introgression between species, as expected, choosing *K* = 3 to represent the gene pools of the three sampled species before gene flow is beneficial for hybridization detection (Fig. 2). However, in cases of hybrid speciation and non-horizontal introgression from a *C*-derived ghost to *B*, opting for *K* = 2, as more readily inferred by the Δ*K* method, proves advantageous for identifying hybridization (Figs. 1c, S3b). In these situations, STRUCTURE with *K* = 3 tends to assign individuals to three distinct pure clusters. It is therefore crucial to go through a range of *K* values, rather than single-mindedly focusing on the “best-*K*” case, if richer, more biologically meaningful insights are to be obtained (Novembre 2016; Toyama et al. 2020).

*Integrating population clustering methods with other techniques to facilitate hybridization detection*—Our results demonstrate that population clustering methods are primarily effective in detecting very recent or ongoing horizontal introgression or hybrid speciation. In these conditions, they can accurately identify hybridization, infer its extent, and determine its direction, even when it involves introgression between sister species. However, the efficacy of population clustering methods is significantly compromised by factors such as post-admixture genetic drift and the presence of ghost lineages, which are prevalent in a phylogenetic context. We have found that, in cases of ancient hybrid speciation, symmetrical contributions from two parental species can erroneously appear highly asymmetrical in admixture analysis using population clustering methods, even completely masking the hybridization signals. Introgression that is not very recent or involves a ghost lineage is often undetectable. These challenges are bolstered by numerous empirical studies documenting cases of hybrid speciation or introgression (Sun et al. 2020; Ayoola et al. 2021; Hirase et al. 2021; Meleshko et al. 2021; Qiao et al. 2021; Wang et al. 2021; Zhang et al. 2024) that were overlooked by population clustering methods yet later confirmed through alternative lines of evidence. Our results strongly suggest that introgression events identified by population clustering methods are likely of a contemporary or recent origin, rather than being attributable to ancient reticulation or ghost introgression.

Additionally, Kong and Kubatko (2021) have warned that multiple hybridization events, which we do not consider in this study, further complicate the admixture analysis. Moreover, spurious admixture signals may be detected by STRUCTURE when there is a large genetic divergence between hybridizing lineages or an excessive amount of ILS or a recent population bottleneck causing strong genetic drift (Lawson et al. 2018; Kong and Kubatko 2021). In summary, population clustering methods alone are often not reliable for detecting hybridization on phylogenetic time-scales.

As a result, we recommend complementing population clustering methods with phylogenetic techniques or more powerful population genetic approaches to infer hybridization events that occur in deep time or originate from ghost lineages. For example, population clustering methods can be initially used to identify recent hybridization and assign individuals into discrete species. This can help exclude admixed individuals from recent hybridization events, ensuring only genetically “pure” individuals to be used in methods for detecting more ancient reticulations or ghost introgression. Phylogenetic network methods that accommodate both ILS and hybridization, such as PhyloNet (Wen et al. 2018) and SNaQ (Solís-Lemus and Ané 2016), are suitable choices for identifying hybridization and introgression in general, despite their limitations in differentiating different types of introgression (Pang and Zhang 2024). More powerful model-based population genetic approaches have been developed and widely applied for the purpose of detecting and quantifying ghost introgression (Ru et al. 2018; Ding et al. 2022; Chang et al. 2023; Pawar et al. 2023; Yamahira et al. 2023; Kato et al. 2024). In this context, full-likelihood methods such as G-PhoCS (Gronau et al. 2011), IMa3 (Hey et al. 2018), and BPP (Flouri et al. 2020; Flouri et al. 2023) hold great promise because they make full use of information in multilocus sequences. Nevertheless, further research and evaluation are needed to examine the performance of these time-consuming population genetic methods in accurately identifying deep reticulation or ghost introgression.

## Conclusion

Population genetic clustering methods are commonly employed for detecting hybridization, even on deep, phylogenetic time-scales. However, our results show that these methods, exemplified by STRUCTURE, struggle to detect signals of hybridization when admixture occurs in the distant past or originates from a ghost lineage. Considering that both ancient hybridization and ghost introgression are much more common over longer time-scales, population genetic clustering methods can significantly underestimate the occurrence of historical or ghost hybridization in empirical systems. Our study points out the significant limitations of population clustering methods in identifying hybridization and underlines the urgent need for developing more efficient and more precise approaches to detect ancient hybridization and ghost introgression events.

## Supplementary Material

Data available from the Dryad Digital Repository:…

## Acknowledgements

We are grateful to Susanne Renner for her encouragement and helpful comments. We would like to thank Ya-Mei Ding, Yang Yang, and Wei-Ning Bai for helpful discussions.

## References

1. Ai H., Fang X., Yang B., Huang Z., Chen H., Mao L., Zhang F., Zhang L., Cui L., He W., Yang J., Yao X., Zhou L., Han L., Li J., Sun S., Xie X., Lai B., Su Y., Lu Y., Yang H., Huang T., Deng W., Nielsen R., Ren J., Huang L. 2015. Adaptation and possible ancient interspecies introgression in pigs identified by whole-genome sequencing. Nat. Genet. 47:217–225.

2. Alexander D. H., Novembre J., Lange K. 2009. Fast model-based estimation of ancestry in unrelated individuals. Genome Res. 19:1655–1664.

3. Anderson E. 1953. Introgressive hybridization. Biol. Rev. Camb. Philos. Soc. 28:280–307.

4. Arnold M. L. 2006. Evolution through genetic exchange. Oxford University Press, USA.

5. Ayoola A. O., Zhang B.-L., Meisel R. P., Nneji L. M., Shao Y., Morenikeji O. B., Adeola A. C., Ng’ang’a S. I., Ogunjemite B. G., Okeyoyin A. O. 2021. Population genomics reveals incipient speciation, introgression, and adaptation in the African mona monkey (*Cercopithecus mona*). Mol. Biol. Evol. 38:876–890.

6. Bai W. N., Yan P. C., Zhang B. W., Woeste K. E., Lin K., Zhang D. Y. 2018. Demographically idiosyncratic responses to climate change and rapid Pleistocene diversification of the walnut genus *Juglans* (Juglandaceae) revealed by whole−genome sequences. New Phytol. 217:1726–1736.

7. Barth J. M., Gubili C., Matschiner M., Tørresen O. K., Watanabe S., Egger B., Han Y.-S., Feunteun E., Sommaruga R., Jehle R. 2020. Stable species boundaries despite ten million years of hybridization in tropical eels. Nat. Commun. 11:1433.

8. Blair C., Ané C. 2020. Phylogenetic trees and networks can serve as powerful and complementary approaches for analysis of genomic data. Syst. Biol. 69:593–601.

9. Blischak P. D., Chifman J., Wolfe A. D., Kubatko L. S. 2018. HyDe: A python package for genome-scale hybridization detection. Syst. Biol. 67:821–829.

10. Bohling J. H., Adams J. R., Waits L. P. 2013. Evaluating the ability of Bayesian clustering methods to detect hybridization and introgression using an empirical red wolf data set. Mol. Ecol. 22:74–86.

11. Chang J. T., Nakamura K., Chao C. T., Luo M. X., Liao P. C. 2023. Ghost introgression facilitates genomic divergence of a sympatric cryptic lineage in *Cycas revoluta*. Ecol. Evol. 13:e10435.

12. Cheng J. Y., Mailund T., Nielsen R. 2017. Fast admixture analysis and population tree estimation for SNP and NGS data. Bioinformatics 33:2148–2155.

13. Degnan J. H. 2018. Modeling hybridization under the network multispecies coalescent. Syst. Biol. 67:786–799.

14. Ding Y.-M., Cao Y., Zhang W.-P., Chen J., Liu J., Li P., Renner S. S., Zhang D.-Y., Bai W.-N. 2022. Population-genomic analyses reveal bottlenecks and asymmetric introgression from Persian into iron walnut during domestication. Genome Biol. 23:1–18.

15. Dittberner H., Tellier A., de Meaux J. 2022. Approximate Bayesian computation untangles signatures of contemporary and historical hybridization between two endangered species. Mol. Biol. Evol. 39: msac015.

16. Edelman N. B., Mallet J. 2021. Prevalence and adaptive impact of introgression. Annu. Rev. Genet. 55:265–283.

17. Evanno G., Regnaut S., Goudet J. 2005. Detecting the number of clusters of individuals using the software STRUCTURE: A simulation study. Mol. Ecol. 14:2611–2620.

18. Flouri T., Jiao X., Huang J., Rannala B., Yang Z. 2023. Efficient Bayesian inference under the multispecies coalescent with migration. Proc. Natl. Acad. Sci. U.S.A. 120:e2310708120.

19. Flouri T., Jiao X., Rannala B., Yang Z. 2020. A Bayesian implementation of the multispecies coalescent model with introgression for phylogenomic analysis. Mol. Biol. Evol. 37:1211–1223.

20. Garcia-Erill G., Albrechtsen A. 2020. Evaluation of model fit of inferred admixture proportions. Mol Ecol Resour 20:936–949.

21. Green R. E., Krause J., Briggs A. W., Maricic T., Stenzel U., Kircher M., Patterson N., Li H., Zhai W., Fritz M. H.-Y. 2010. A draft sequence of the Neandertal genome. Science 328:710–722.

22. Gronau I., Hubisz M. J., Gulko B., Danko C. G., Siepel A. 2011. Bayesian inference of ancient human demography from individual genome sequences. Nat. Genet. 43:1031– 1034.

23. Hey J., Chung Y., Sethuraman A., Lachance J., Tishkoff S., Sousa V. C., Wang Y. 2018. Phylogeny estimation by integration over isolation with migration models. Mol. Biol. Evol. 35:2805–2818.

24. Hey J., Nielsen R. 2004. Multilocus methods for estimating population sizes, migration rates and divergence time, with applications to the divergence of *Drosophila pseudoobscura* and *D. persimilis*. Genetics 167:747–760.

25. Hibbins M. S., Hahn M. W. 2019. The timing and direction of introgression under the multispecies network coalescent. Genetics 211:1059–1073.

26. Hibbins M. S., Hahn M. W. 2022. Phylogenomic approaches to detecting and characterizing introgression. Genetics 220:iyab173.

27. Hirase S., Yamasaki Y. Y., Sekino M., Nishisako M., Ikeda M., Hara M., Merilä J., Kikuchi K. 2021. Genomic evidence for speciation with gene flow in broadcast spawning marine invertebrates. Mol. Biol. Evol. 38:4683–4699.

28. Hudson R. R. 2002. Generating samples under a wright-fisher neutral model of genetic variation. Bioinformatics 18:337–338.

29. Janes J. K., Miller J. M., Dupuis J. R., Malenfant R. M., Gorrell J. C., Cullingham C. I., Andrew R. L. 2017. The k = 2 conundrum. Mol. Ecol. 26:3594–3602.

30. Jukes T. H., Cantor C. R. 1969. Evolution of protein molecules. Mammalian protein metabolism. New York: Academic Press, p. 21–132.

31. Kato S., Arakaki S., Nagano A. J., Kikuchi K., Hirase S. 2024. Genomic landscape of introgression from the ghost lineage in a gobiid fish uncovers the generality of forces shaping hybrid genomes. Mol. Ecol. doi: 10.1111/mec.17216.

32. Kong S., Kubatko L. S. 2021. Comparative performance of popular methods for hybrid detection using genomic data. Syst. Biol. 70:891–907.

33. Kuhlwilm M., Han S., Sousa V. C., Excoffier L., Marques-Bonet T. 2019. Ancient admixture from an extinct ape lineage into bonobos. Nat. Ecol. Evol. 3:957–965.

34. Lawson D. J., van Dorp L., Falush D. 2018. A tutorial on how not to over-interpret STRUCTURE and ADMIXTURE bar plots. Nat. Commun. 9:3258.

35. Li J., Milne R. I., Ru D., Miao J., Tao W., Zhang L., Xu J., Liu J., Mao K. 2020. Allopatric divergence and hybridization within *Cupressus chengiana* (Cupressaceae), a threatened conifer in the northern hengduan mountains of western china. Mol. Ecol. 29:1250–1266.

36. Li J., Zhang Y., Ruhsam M., Milne R. I., Wang Y., Wu D., Jia S., Tao T., Mao K. 2022. Seeing through the hedge: Phylogenomics of *Thuja* (Cupressaceae) reveals prominent incomplete lineage sorting and ancient introgression for Tertiary relict flora. Cladistics 38:187–203.

37. Li M., Zheng Z., Liu J., Yang Y., Ren G., Ru D., Zhang S., Du X., Ma T., Milne R., Liu J. 2021. Evolutionary origin of a tetraploid *Allium* species on the Qinghai–Tibet Plateau. Mol. Ecol. 30:5780–5795.

38. Li X., Wei G., El-Kassaby Y. A., Fang Y. 2021. Hybridization and introgression in sympatric and allopatric populations of four oak species. BMC Plant Biol. 21:266.

39. Lopes F., Oliveira L. R., Beux Y., Kessler A., Cárdenas-Alayza S., Majluf P., Páez-Rosas D., Chaves J., Crespo E., Brownell Jr R. L. 2023. Genomic evidence for homoploid hybrid speciation in a marine mammal apex predator. Sci. Adv. 9:eadf6601.

40. Mallet J. 2005. Hybridization as an invasion of the genome. Trends Ecol. Evol. 20:229–237.

41. Mallet J. 2007. Hybrid speciation. Nature 446:279–283.

42. Mallet J., Besansky N., Hahn M. W. 2016. How reticulated are species? Bioessays 38:140– 149.

43. Meleshko O., Martin M. D., Korneliussen T. S., Schröck C., Lamkowski P., Schmutz J., Healey A., Piatkowski B. T., Shaw A. J., Weston D. J., Flatberg K. I., Szövényi P., Hassel K., Stenøien H. K. 2021. Extensive genome-wide phylogenetic discordance is due to incomplete lineage sorting and not ongoing introgression in a rapidly radiated bryophyte genus. Mol. Biol. Evol. 38:2750–2766.

44. Neophytou C. 2014. Bayesian clustering analyses for genetic assignment and study of hybridization in oaks: Effects of asymmetric phylogenies and asymmetric sampling schemes. Tree Genet. Genom. 10:273–285.

45. Novembre J. 2016. Pritchard, stephens, and donnelly on population structure. Genetics 204:391–393.

46. Oliveira R., Randi E., Mattucci F., Kurushima J., Lyons L., Alves P. 2015. Toward a genome-wide approach for detecting hybrids: Informative snps to detect introgression between domestic cats and European wildcats (*Felis silvestris*). Heredity 115:195–205.

47. Ottenburghs J. 2020. Ghost introgression: Spooky gene flow in the distant past. Bioessays 42:e2000012.

48. Pang X.-X., Zhang D.-Y. 2023. Impact of ghost introgression on coalescent-based species tree inference and estimation of divergence time. Syst. Biol. 72:35–49.

49. Pang X.-X., Zhang D.-Y. 2024. Detection of ghost introgression requires exploiting topological and branch length information. Syst. Biol.:10.1093/sysbio/syad1077.

50. Pawar H., Rymbekova A., Cuadros-Espinoza S., Huang X., de Manuel M., van der Valk T., Lobon I., Alvarez-Estape M., Haber M., Dolgova O., Han S., Esteller-Cucala P., Juan D., Ayub Q., Bautista R., Kelley J. L., Cornejo O. E., Lao O., Andrés A. M., Guschanski K., Ssebide B., Cranfield M., Tyler-Smith C., Xue Y., Prado-Martinez J., Marques-Bonet T., Kuhlwilm M. 2023. Ghost admixture in eastern gorillas. Nat. Ecol. Evol. 7:1503–1514.

51. Pritchard J. K., Stephens M., Donnelly P. 2000. Inference of population structure using multilocus genotype data. Genetics 155:945–959.

52. Qiao Q., Edger P. P., Xue L., Qiong L., Lu J., Zhang Y., Cao Q., Yocca A. E., Platts A. E., Knapp S. J. 2021. Evolutionary history and pan-genome dynamics of strawberry (*Fragaria* spp.). Proc. Natl. Acad. Sci. U.S.A. 118:e2105431118.

53. Rannala B., Yang Z. 2017. Efficient Bayesian species tree inference under the multispecies coalescent. Syst. Biol. 66:823–842.

54. Ravagni S., Sanchez−Donoso I., Vil◊ C. 2021. Biased assessment of ongoing admixture using structure in the absence of reference samples. Mol Ecol Resour 21:677–689.

55. Rocha J. L., Vaz Pinto P., Siegismund H. R., Meyer M., Jansen van Vuuren B., Veríssimo L., Ferrand N., Godinho R. 2022. African climate and geomorphology drive evolution and ghost introgression in sable antelope. Mol. Ecol. 31:2968–2984.

56. Ru D., Sun Y., Wang D., Chen Y., Wang T., Hu Q., Abbott R. J., Liu J. 2018. Population genomic analysis reveals that homoploid hybrid speciation can be a lengthy process. Mol. Ecol. 27:4875–4887.

57. Sankararaman S., Mallick S., Dannemann M., Prüfer K., Kelso J., Pääbo S., Patterson N., Reich D. 2014. The genomic landscape of Neanderthal ancestry in present-day humans. Nature 507:354–357.

58. Schumer M., Rosenthal G. G., Andolfatto P. 2014. How common is homoploid hybrid speciation? Evolution 68:1553–1560.

59. Schwenk K., Brede N., Streit B. 2008. Introduction. Extent, processes and evolutionary impact of interspecific hybridization in animals. Philos. Trans. R. Soc. Lond., B, Biol. Sci. 363:2805–2811.

60. Solís-Lemus C., Ané C. 2016. Inferring phylogenetic networks with maximum pseudolikelihood under incomplete lineage sorting. PLoS Genet. 12:e1005896.

61. Sørensen E. F., Harris R. A., Zhang L., Raveendran M., Kuderna L. F. K., Walker J. A., Storer J. M., Kuhlwilm M., Fontsere C., Seshadri L., Bergey C. M., Burrell A. S., Bergman J., Phillips-Conroy J. E., Shiferaw F., Chiou K. L., Chuma I. S., Keyyu J. D., Fischer J., Gingras M.-C., Salvi S., Doddapaneni H., Schierup M. H., Batzer M. A., Jolly C. J., Knauf S., Zinner D., Farh K. K.-H., Marques-Bonet T., Munch K., Roos C., Rogers J. 2023. Genome-wide coancestry reveals details of ancient and recent male-driven reticulation in baboons. Science 380:eabn8153.

62. Stull G. W., Pham K. K., Soltis P. S., Soltis D. E. 2023. Deep reticulation: The long legacy of hybridization in vascular plant evolution. Plant J. 114:743–766.

63. Sun Y., Lu Z., Zhu X., Ma H. 2020. Genomic basis of homoploid hybrid speciation within chestnut trees. Nat. Commun. 11:3375.

64. Taylor S. A., Larson E. L. 2019. Insights from genomes into the evolutionary importance and prevalence of hybridization in nature. Nat. Ecol. Evol. 3:170–177.

65. Thawornwattana Y., Huang J., Flouri T., Mallet J., Yang Z. 2023. Inferring the direction of introgression using genomic sequence data. Mol. Biol. Evol. 40:msad178.

66. Tiley G. P., Flouri T., Jiao X., Poelstra J. W., Xu B., Zhu T., Rannala B., Yoder A. D., Yang Z. 2023. Estimation of species divergence times in presence of cross-species gene flow. Syst. Biol. 72:820–836.

67. Toyama K. S., Crochet P. A., Leblois R. 2020. Sampling schemes and drift can bias admixture proportions inferred by structure. Mol Ecol Resour 20:1769–1785.

68. Tricou T., Tannier E., de Vienne D. M. 2022a. Ghost lineages highly influence the interpretation of introgression tests. Syst. Biol. 71:1147–1158.

69. Tricou T., Tannier E., de Vienne D. M. 2022b. Ghost lineages can invalidate or even reverse findings regarding gene flow. PLoS Biol. 20:e3001776.

70. van Wyk A. M., Dalton D. L., Hoban S., Bruford M. W., Russo I. R. M., Birss C., Grobler P., van Vuuren B. J., Kotzé A. 2017. Quantitative evaluation of hybridization and the impact on biodiversity conservation. Ecol. Evol. 7:320–330.

71. Wang J. 2019. A parsimony estimator of the number of populations from a STRUCTURE− like analysis. Mol Ecol Resour 19:970–981.

72. Wang M.-S., Wang S., Li Y., Jhala Y., Thakur M., Otecko N. O., Si J.-F., Chen H.-M., Shapiro B., Nielsen R. 2020. Ancient hybridization with an unknown population facilitated high-altitude adaptation of canids. Mol. Biol. Evol. 37:2616–2629.

73. Wang Z., Jiang Y., Bi H., Lu Z., Ma Y., Yang X., Chen N., Tian B., Liu B., Mao X. 2021. Hybrid speciation via inheritance of alternate alleles of parental isolating genes. Mol. Plant 14:208–222.

74. Wen D., Yu Y., Zhu J., Nakhleh L. 2018. Inferring phylogenetic networks using PhyloNet. Syst. Biol. 67:735–740.

75. White O. W., Reyes-Betancort A., Chapman M. A., Carine M. A. 2018. Independent homoploid hybrid speciation events in the Macaronesian endemic genus *Argyranthemum*. Mol. Ecol. 27:4856–4874.

76. Whitney K. D., Ahern J. R., Campbell L. G., Albert L. P., King M. S. 2010. Patterns of hybridization in plants. Perspect. Plant Ecol. Evol. Syst. 12:175–182.

77. Wu H., Wang Z., Zhang Y., Frantz L., Roos C., Irwin D. M., Zhang C., Liu X., Wu D., Huang S. 2023. Hybrid origin of a primate, the gray snub-nosed monkey. Science 380:eabl4997.

78. Yamahira K., Kobayashi H., Kakioka R., Montenegro J., Masengi K. W., Okuda N., Nagano A. J., Tanaka R., Naruse K., Tatsumoto S. 2023. Ghost introgression in ricefishes of the genus *Adrianichthys* in an ancient Wallacean lake. J. Evol. Biol. 36:1484–1493.

79. Yang Z., Flouri T. 2022. Estimation of cross-species introgression rates using genomic data despite model unidentifiability. Mol. Biol. Evol. 39:msac083.

80. Yu J., Niu Y., You Y., Cox C. J., Barrett R. L., Trias−Blasi A., Guo J., Wen J., Lu L., Chen Z. 2023. Integrated phylogenomic analyses unveil reticulate evolution in *Parthenocissus* (Vitaceae), highlighting speciation dynamics in the Himalayan–Hengduan mountains. New Phytol. 238:888–903.

81. Yu Y., Dong J., Liu K. J., Nakhleh L. 2014. Maximum likelihood inference of reticulate evolutionary histories. Proc. Natl. Acad. Sci. USA 111:16448–16453.

82. Zhang B.-W., Xu L.-L., Li N., Yan P.-C., Jiang X.-H., Woeste K. E., Lin K., Renner S. S., Zhang D.-Y., Bai W.-N. 2019. Phylogenomics reveals an ancient hybrid origin of the Persian walnut. Mol. Biol. Evol. 36:2451–2461.

83. Zhang D., She H., Wang S., Wang H., Li S., Cheng Y., Song G., Jia C., Qu Y., Rheindt F. E. 2024. Phylogenetic conflict between species tree and maternally inherited gene trees in a clade of *Emberiza* buntings (aves: Emberizidae). Syst. Biol.:syad078.

84. Zheng Y., Janke A. 2018. Gene flow analysis method, the D-statistic, is robust in a wide parameter space. BMC Bioinform. 19:10.

85. Zou T., Kuang W., Yin T., Frantz L., Zhang C., Liu J., Wu H., Yu L. 2022. Uncovering the enigmatic evolution of bears in greater depth: The hybrid origin of the Asiatic black bear. Proc. Natl. Acad. Sci. U.S.A. 119:e2120307119.

