## supplemental figures for "A Cautionary Note on Using STRUCTURE to Detect Hybridization in a Phylogenetic Context"

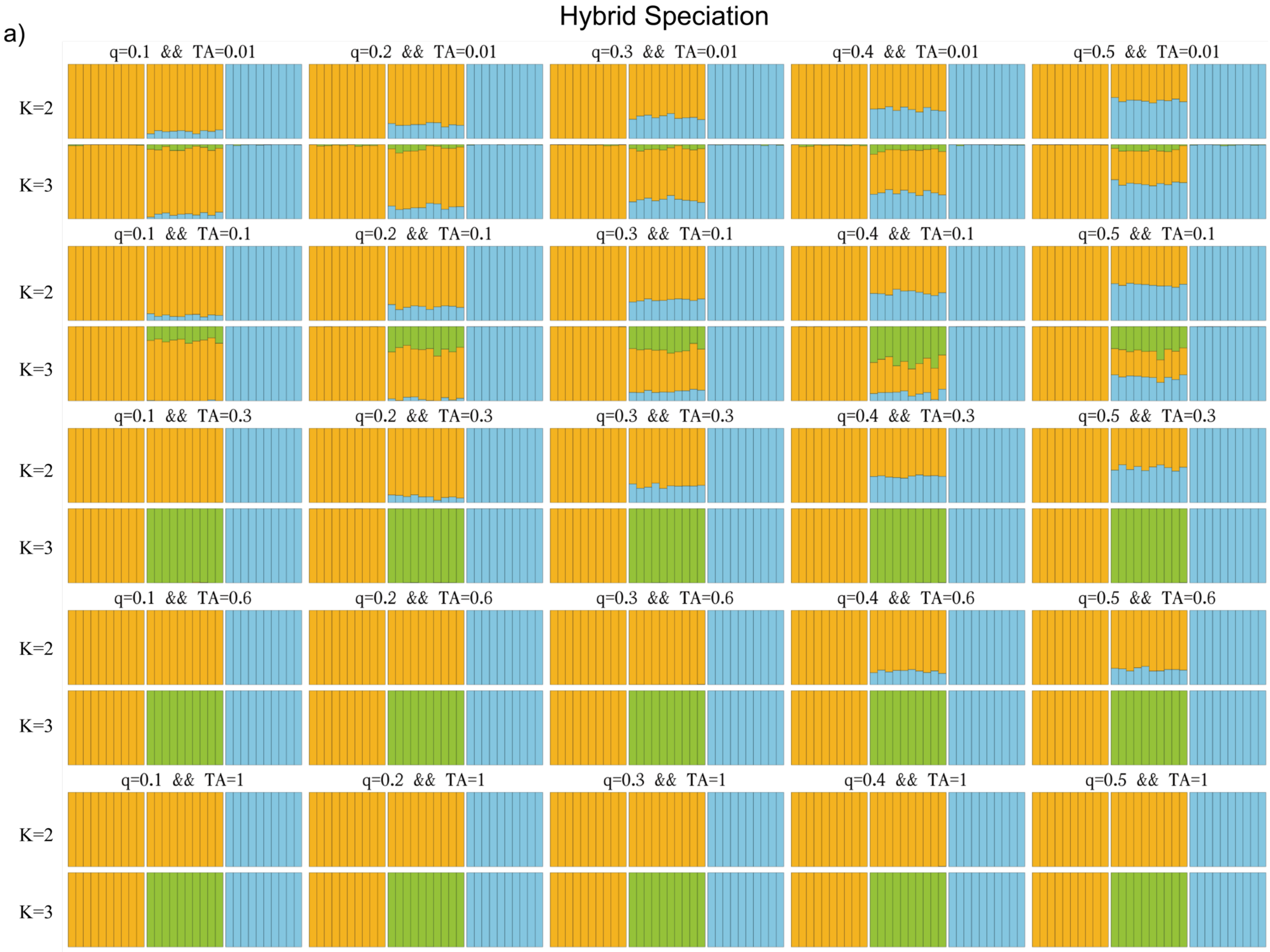
a)
Hybrid Speciation
q=0.1 && TA=0.01
q=0.2 && TA=0.01
q=0.3 && TA=0.01
q=0.4 && TA=0.01
q=0.5 && TA=0.01
q=0.1 && TA=0.1
q=0.2 && TA=0.1
q=0.3 && TA=0.1
q=0.4 && TA=0.1
q=0.5 && TA=0.1
q=0.1 && TA=0.3
q=0.2 && TA=0.3
q=0.3 && TA=0.3
q=0.4 && TA=0.3
q=0.5 && TA=0.3
q=0.1 && TA=0.6
q=0.2 && TA=0.6
q=0.3 && TA=0.6
q=0.4 && TA=0.6
q=0.5 && TA=0.6
q=0.1 && TA=1
q=0.2 && TA=1
q=0.3 && TA=1
q=0.4 && TA=1
q=0.5 && TA=1
K=2
K=3

1

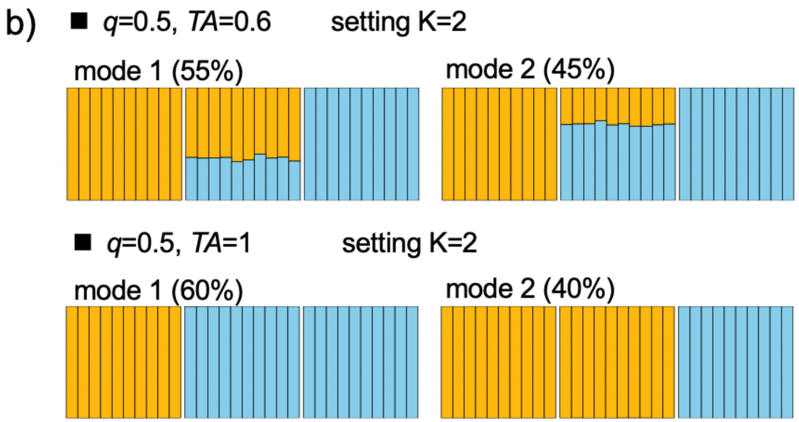


Figure S1. Inferred STRUCTURE clustering plots for *K* = 2 and *K* = 3 under various combinations of parameters in the scenarios of hybrid speciation (Fig. 1a). a) The major clustering mode for individuals from species *A*, *B*, and *C* (from left to right). The top of each plot indicates the proportion of genetic material in hybrids inherited from parent *C* (*q*) and the time of the admixture event (*TA*). b) Cases of multiple distinct clustering modes, each supported by at least 20% of the replicates.

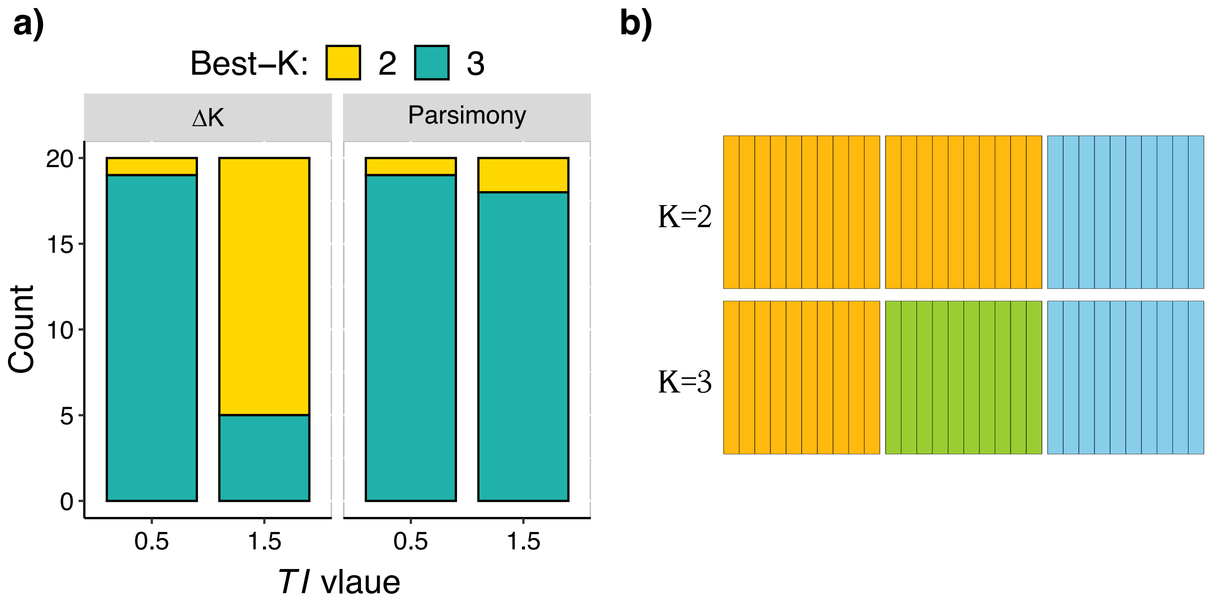


Figure S2. Results for scenarios of species tree *AB*|*C* in the absence of introgression (i.e., $q$ = 0) in Figure 2a. a) Best-*K* inference using Δ*K* and parsimony methods. Colored bars represent the numbers of inferences for best-$K$ = 2 and best-$K$ = 3 among 20 replicates. The strip at the top of each plot indicates the methods employed. The x-axis indicates the values of *TI*. b) Inferred STRUCTURE clustering plots when specifying $K$ = 2 or $K$ = 3. Each plot illustrates the mode of clustering for individuals from species *A*, *B*, and *C* (from left to right). The clustering results remain consistent across 20 replicates and *TI* values.

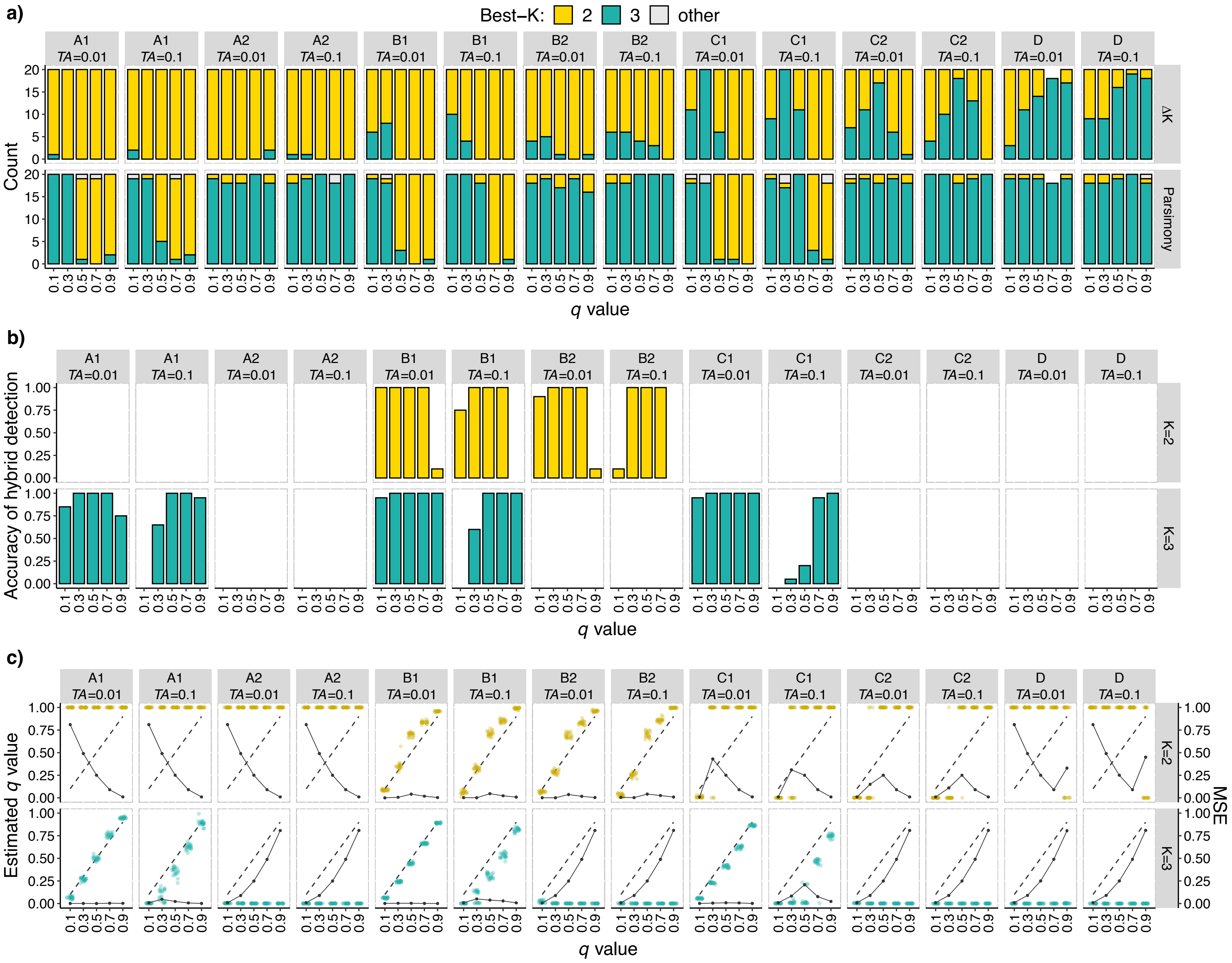

a)
Best−K: 2 3 other
A1 TA=0.01
A1 TA=0.1
A2 TA=0.01
A2 TA=0.1
B1 TA=0.01
B1 TA=0.1
B2 TA=0.01
B2 TA=0.1
C1 TA=0.01
C1 TA=0.1
C2 TA=0.01
C2 TA=0.1
D TA=0.01
D TA=0.1
Count
ΔK
Parsimony
q value
b)
Accuracy of hybrid detection
K=2
K=3
q value
c)
Estimated q value
MSE
K=2
K=3
q value

Figure S3. Results for scenarios of unidirectional horizontal or non-horizontal (ghost) introgression with the parameter $TI = 1.5$ (Fig. 2a). a) Best-$K$ inference using $\Delta K$ and parsimony methods. Colored bars represent the numbers of inferences for best-$K$ = 2 and best-$K$ = 3 among 20 replicates. The strip at the top of each plot indicates the simulated scenarios and the parameter $TA$, while the strip on the right of each plot represents the methods employed. The x-axis indicates the values of $q$. b-c) Plots of the results for hybrid detection and estimation of admixture proportion ($q$-value) assuming $K = 2$ and $K = 3$. The strip at the top of each plot indicates the simulated scenarios and the parameter $TA$. The strips on the right of each plot represent the chosen number of clusters $K$. The x-axis indicates the values of $q$. b) Accuracy of hybrid detection. The y-axis represents the proportion of times that hybrid individuals were successfully identified. c) Estimation of admixture proportion $q$-value. Colored points represent estimates of $q$, horizontally jittered to avoid clutter. The dashed line shows the true values, while the solid line with points represents the MSE values.

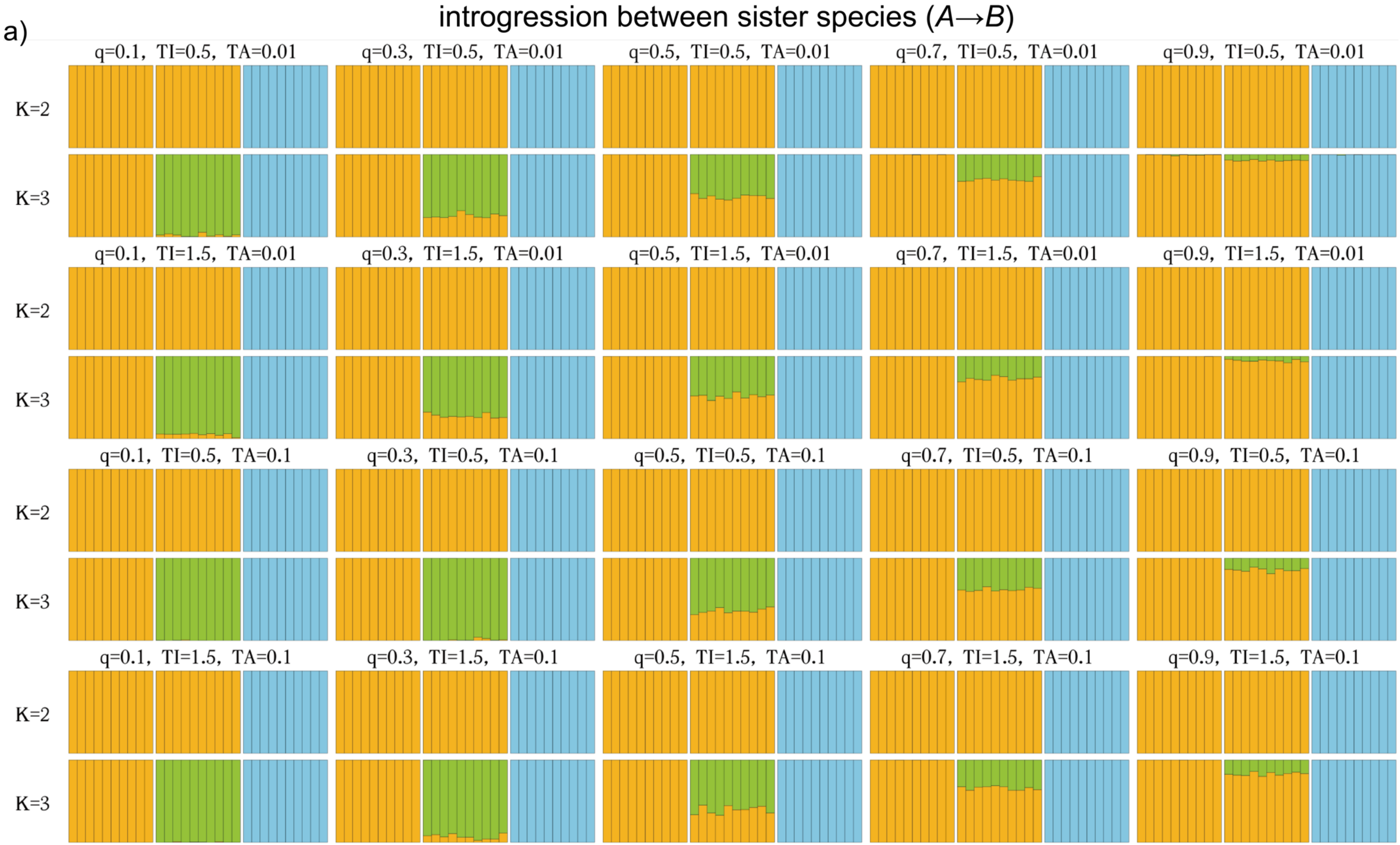
a)
introgression between sister species (A→B)
q=0.1, TI=0.5, TA=0.01
q=0.3, TI=0.5, TA=0.01
q=0.5, TI=0.5, TA=0.01
q=0.7, TI=0.5, TA=0.01
q=0.9, TI=0.5, TA=0.01
q=0.1, TI=1.5, TA=0.01
q=0.3, TI=1.5, TA=0.01
q=0.5, TI=1.5, TA=0.01
q=0.7, TI=1.5, TA=0.01
q=0.9, TI=1.5, TA=0.01
q=0.1, TI=0.5, TA=0.1
q=0.3, TI=0.5, TA=0.1
q=0.5, TI=0.5, TA=0.1
q=0.7, TI=0.5, TA=0.1
q=0.9, TI=0.5, TA=0.1
q=0.1, TI=1.5, TA=0.1
q=0.3, TI=1.5, TA=0.1
q=0.5, TI=1.5, TA=0.1
q=0.7, TI=1.5, TA=0.1
q=0.9, TI=1.5, TA=0.1
K=2
K=3

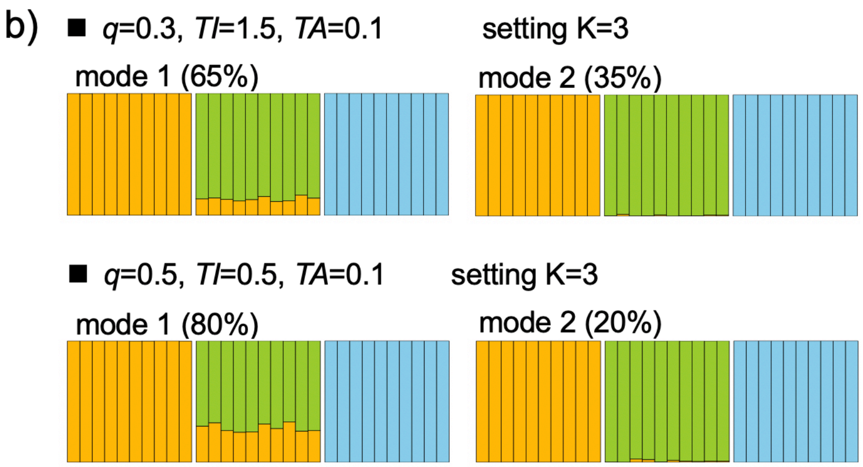


Figure S4. Inferred STRUCTURE clustering plots for $K = 2$ and $K = 3$ under various combinations of parameters in the scenarios of introgression between sister species (Scenario A1 in Fig. 2a). a) The major clustering mode for individuals from species *A*, *B*, and *C* (from left to right). The top of each plot indicates the proportion of genetic material in hybrids inherited from parent *C* (*q*), the time interval between two speciation events (*TI*), and the timing of admixture (*TA*). b) Cases of multiple distinct clustering modes, each supported by at least 20% of the replicates.

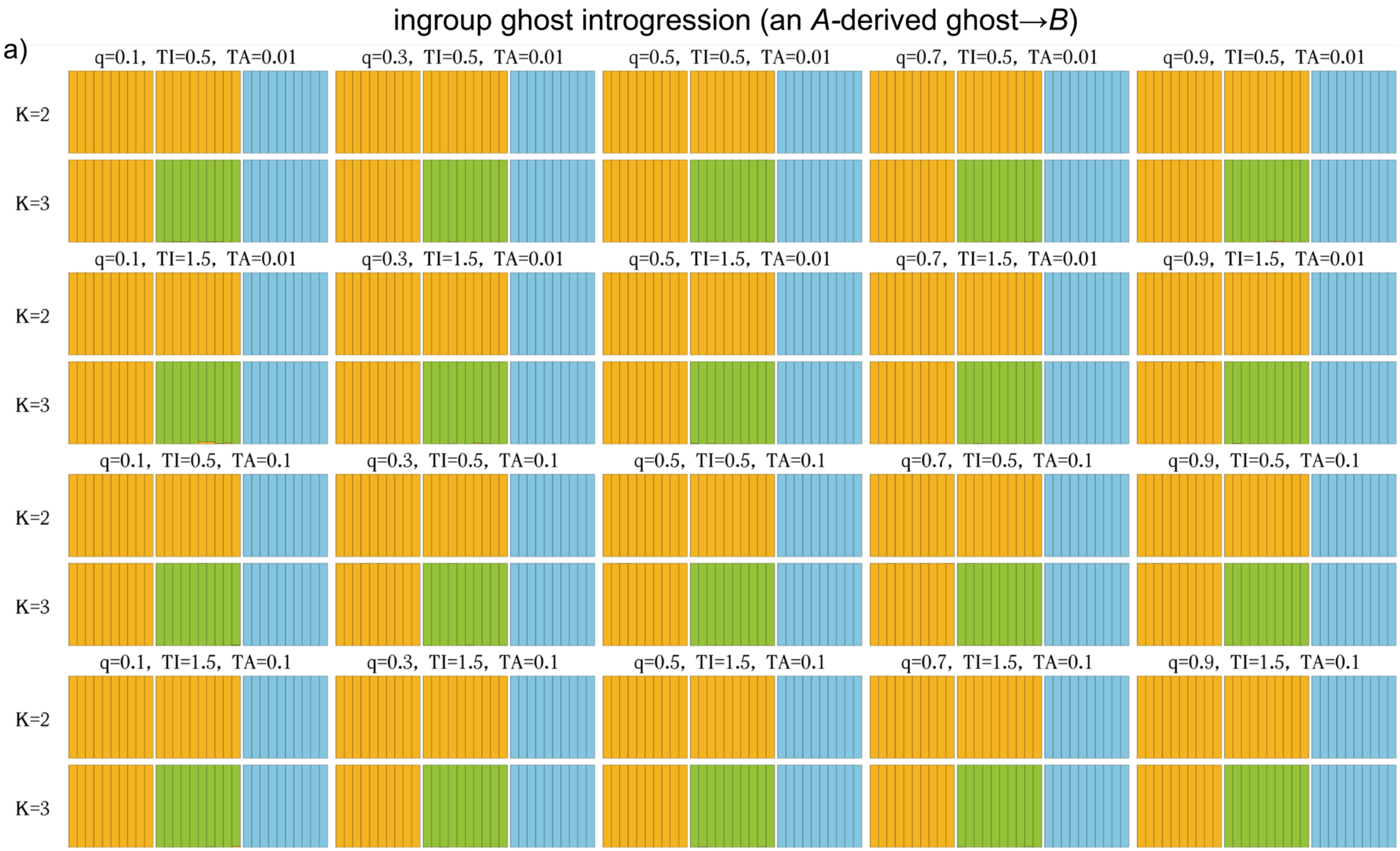
ingroup ghost introgression (an A-derived ghost→B)
a)
q=0.1, TI=0.5, TA=0.01
q=0.3, TI=0.5, TA=0.01
q=0.5, TI=0.5, TA=0.01
q=0.7, TI=0.5, TA=0.01
q=0.9, TI=0.5, TA=0.01
K=2
K=3
q=0.1, TI=1.5, TA=0.01
q=0.3, TI=1.5, TA=0.01
q=0.5, TI=1.5, TA=0.01
q=0.7, TI=1.5, TA=0.01
q=0.9, TI=1.5, TA=0.01
K=2
K=3
q=0.1, TI=0.5, TA=0.1
q=0.3, TI=0.5, TA=0.1
q=0.5, TI=0.5, TA=0.1
q=0.7, TI=0.5, TA=0.1
q=0.9, TI=0.5, TA=0.1
K=2
K=3
q=0.1, TI=1.5, TA=0.1
q=0.3, TI=1.5, TA=0.1
q=0.5, TI=1.5, TA=0.1
q=0.7, TI=1.5, TA=0.1
q=0.9, TI=1.5, TA=0.1
K=2
K=3

Figure S5. Inferred STRUCTURE clustering plots for $K = 2$ and $K = 3$ under various combinations of parameters in the scenarios of introgression from an *A*-derived ghost to *B* (Scenario A2 in Fig. 2a). At the top of each plot, the proportion of genetic material in hybrids inherited from the donor (*q*), the time interval between two speciation events (*TI*), and the timing of admixture (*TA*) are indicated. Each plot illustrates the major mode of clustering for individuals from species *A*, *B*, and *C* (from left to right) when assuming $K = 2$ or $K = 3$. No case encompasses multiple distinct clustering modes.

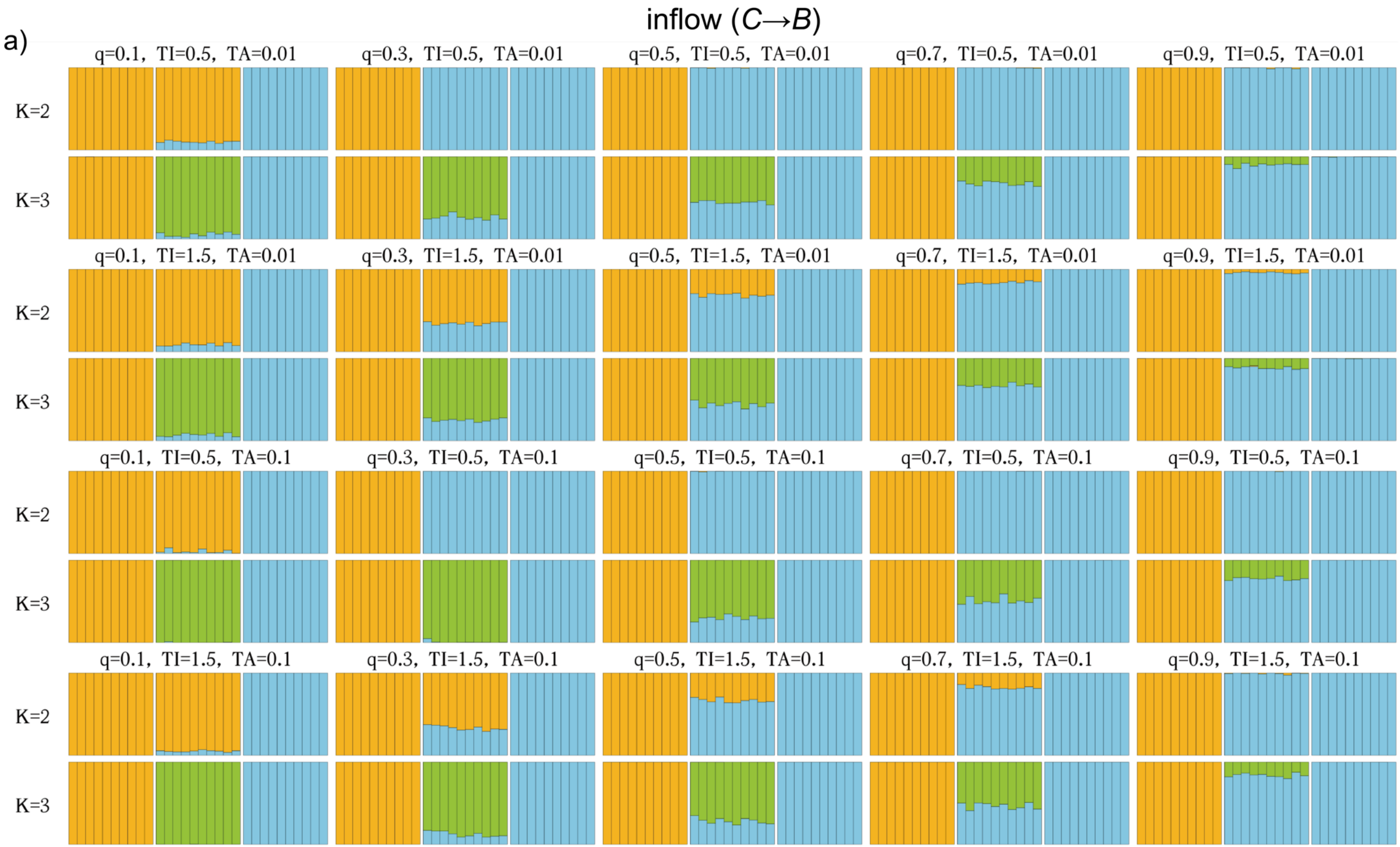
inflow (C→B)
a)
q=0.1, TI=0.5, TA=0.01
q=0.3, TI=0.5, TA=0.01
q=0.5, TI=0.5, TA=0.01
q=0.7, TI=0.5, TA=0.01
q=0.9, TI=0.5, TA=0.01
q=0.1, TI=1.5, TA=0.01
q=0.3, TI=1.5, TA=0.01
q=0.5, TI=1.5, TA=0.01
q=0.7, TI=1.5, TA=0.01
q=0.9, TI=1.5, TA=0.01
q=0.1, TI=0.5, TA=0.1
q=0.3, TI=0.5, TA=0.1
q=0.5, TI=0.5, TA=0.1
q=0.7, TI=0.5, TA=0.1
q=0.9, TI=0.5, TA=0.1
q=0.1, TI=1.5, TA=0.1
q=0.3, TI=1.5, TA=0.1
q=0.5, TI=1.5, TA=0.1
q=0.7, TI=1.5, TA=0.1
q=0.9, TI=1.5, TA=0.1
K=2
K=3

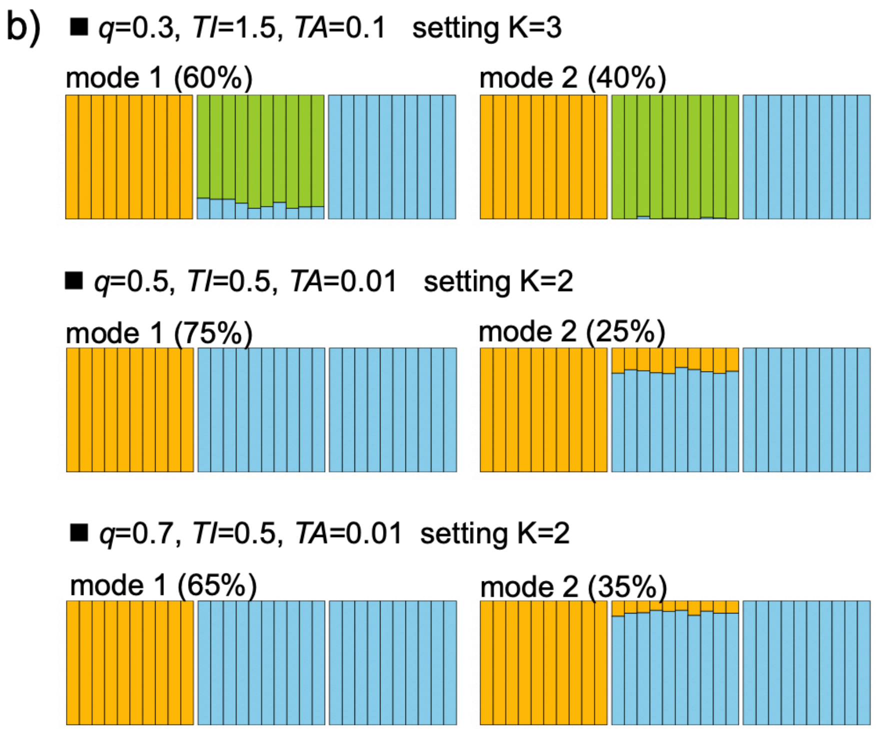


Figure S6. Inferred STRUCTURE clustering plots for $K = 2$ and $K = 3$ under various combinations of parameters in the scenarios of inflow introgression (Scenario B1 in Fig. 2a). a) The major clustering mode for individuals from species *A*, *B*, and *C* (from left to right). The top of each plot indicates the proportion of genetic material in hybrids inherited from parent *C* ($q$), the time interval between two speciation events (*TI*), and the timing of admixture (*TA*). b) Cases of multiple distinct clustering modes, each supported by at least 20% of the replicates.

ingroup ghost introgression (a *C*-derived ghost→*B*)

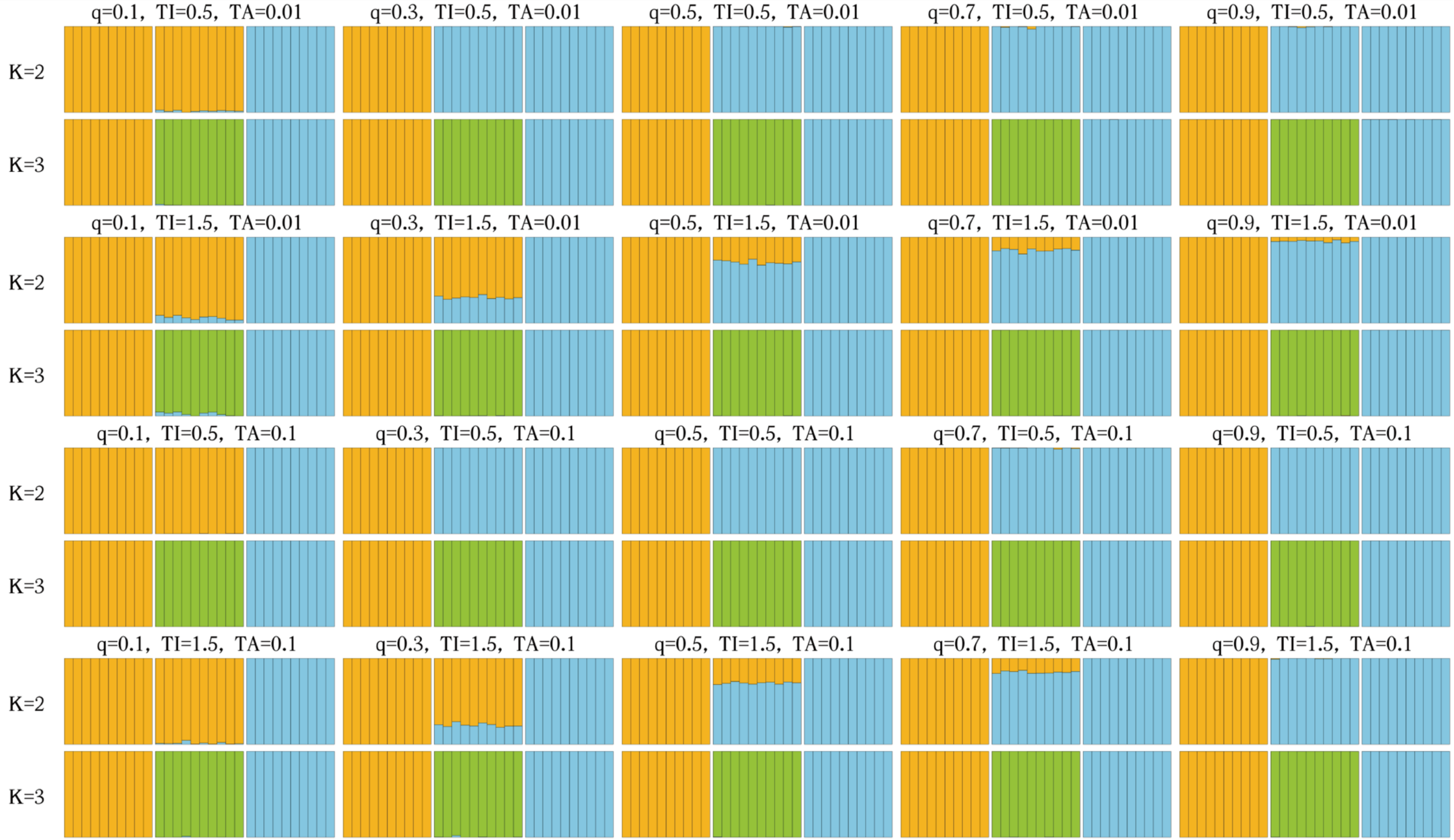

Figure S7. Inferred STRUCTURE clustering plots for $K = 2$ and $K = 3$ under various combinations of parameters in the scenarios of introgression from a *C*-derived ghost to *B* (Scenario B2 in Fig. 2a). At the top of each plot, the proportion of genetic material in hybrids inherited from the donor ($q$), the time interval between two speciation events ($TI$), and the timing of admixture ($TA$) are indicated. Each plot illustrates the major mode of clustering for individuals from species *A*, *B*, and *C* (from left to right) when assuming $K = 2$ or $K = 3$. No case encompasses multiple distinct clustering modes.

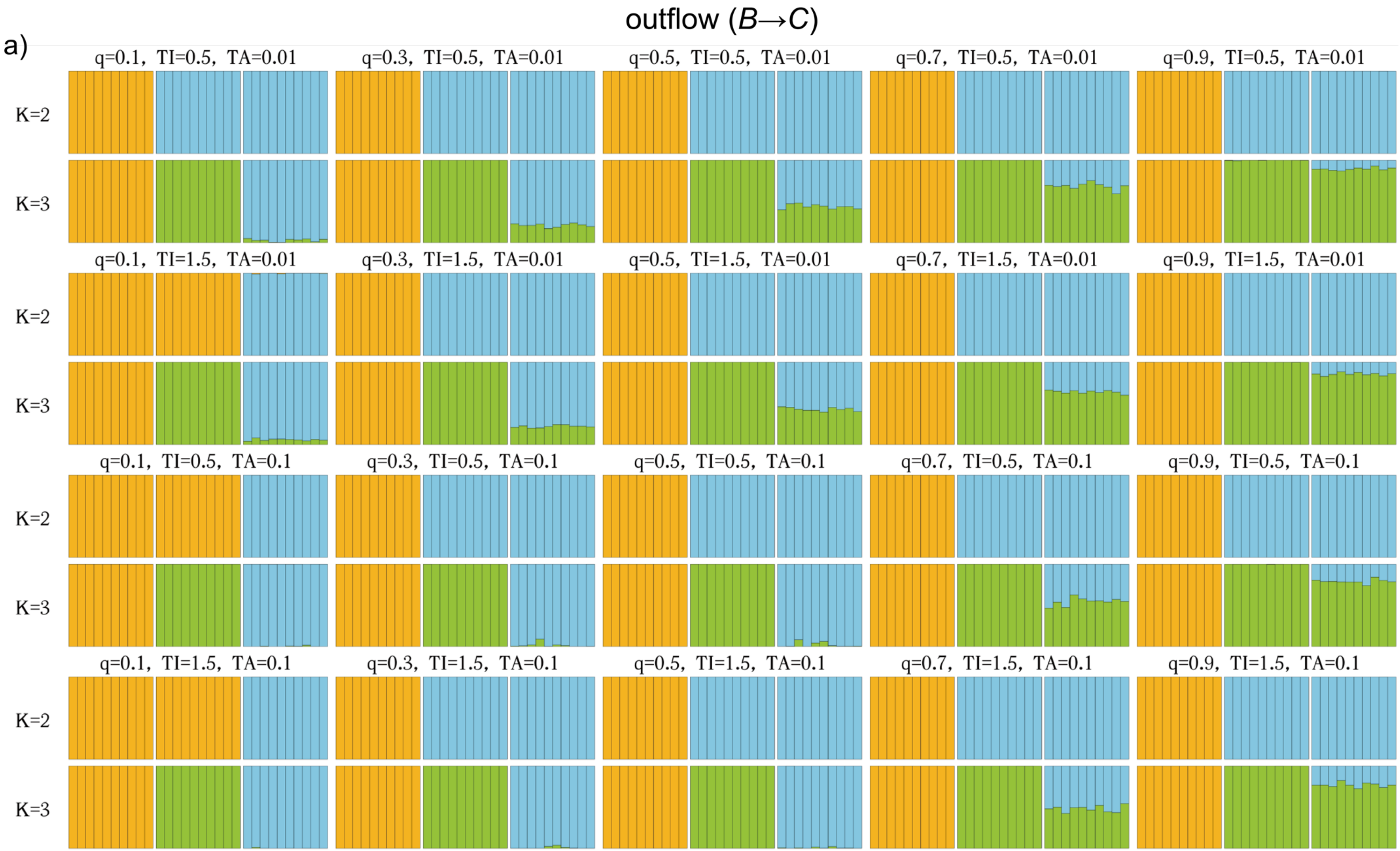
outflow (B→C)
a)
q=0.1, TI=0.5, TA=0.01
q=0.3, TI=0.5, TA=0.01
q=0.5, TI=0.5, TA=0.01
q=0.7, TI=0.5, TA=0.01
q=0.9, TI=0.5, TA=0.01
K=2
K=3
q=0.1, TI=1.5, TA=0.01
q=0.3, TI=1.5, TA=0.01
q=0.5, TI=1.5, TA=0.01
q=0.7, TI=1.5, TA=0.01
q=0.9, TI=1.5, TA=0.01
K=2
K=3
q=0.1, TI=0.5, TA=0.1
q=0.3, TI=0.5, TA=0.1
q=0.5, TI=0.5, TA=0.1
q=0.7, TI=0.5, TA=0.1
q=0.9, TI=0.5, TA=0.1
K=2
K=3
q=0.1, TI=1.5, TA=0.1
q=0.3, TI=1.5, TA=0.1
q=0.5, TI=1.5, TA=0.1
q=0.7, TI=1.5, TA=0.1
q=0.9, TI=1.5, TA=0.1
K=2
K=3

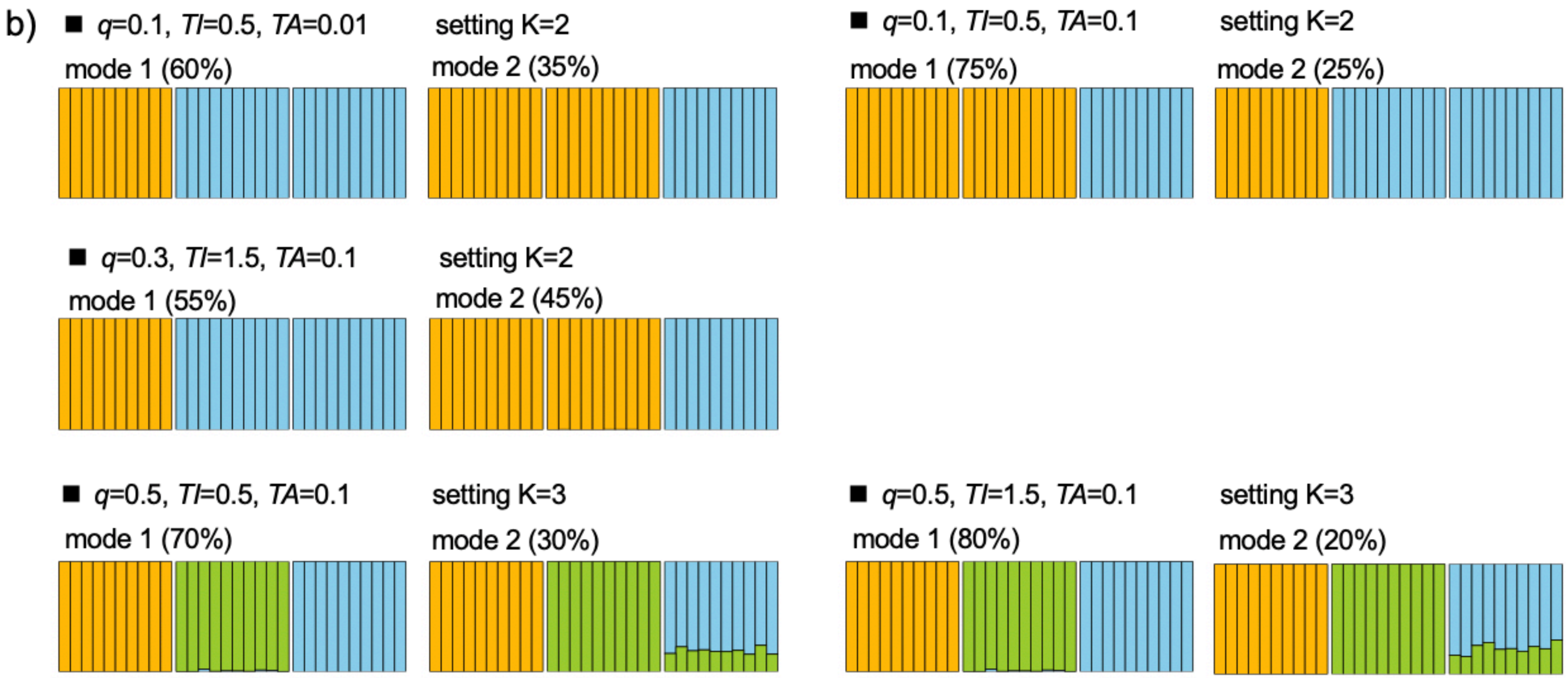


Figure S8. Inferred STRUCTURE clustering plots for $K = 2$ and $K = 3$ under various combinations of parameters in the scenarios of outflow introgression (Scenario C1 in Fig. 2a). a) The major clustering mode for individuals from species *A*, *B*, and *C* (from left to right). The top of each plot indicates the proportion of genetic material in hybrids inherited from parent *C* ($q$), the time interval between two speciation events (*TI*), and the timing of admixture (*TA*). b) Cases of multiple distinct clustering modes, each supported by at least 20% of the replicates.

ingroup ghost introgression (a *B*-derived ghost→*C*)

a)

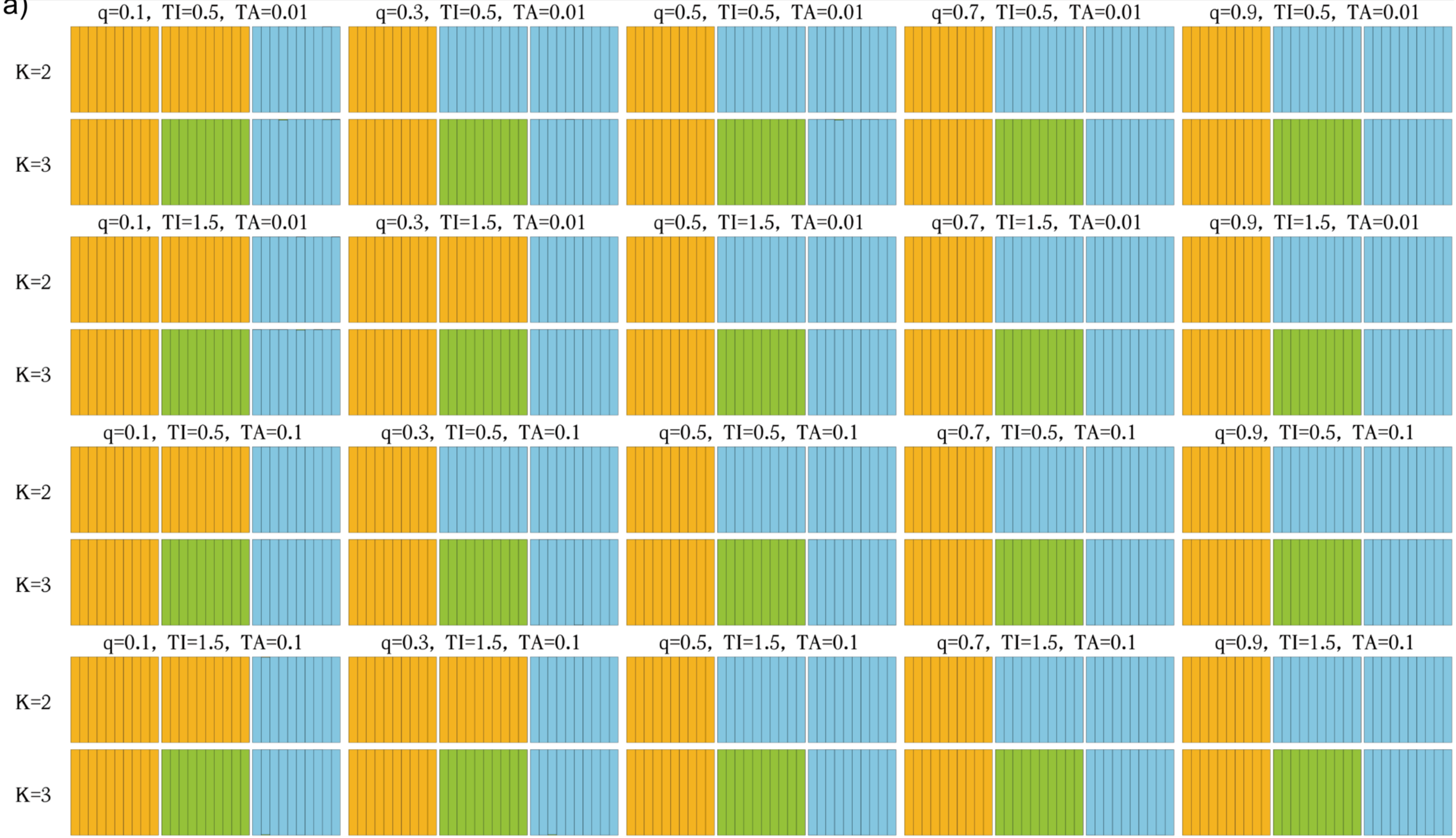


55

Figure S9. Inferred STRUCTURE clustering plots for $K = 2$ and $K = 3$ under various combinations of parameters in the scenarios of introgression from a *B*-derived ghost to *C* (Scenario C2 in Fig. 2a). At the top of each plot, the proportion of genetic material in hybrids inherited from the donor (*q*), the time interval between two speciation events (*TI*), and the timing of admixture (*TA*) are indicated. Each plot illustrates the major mode of clustering for individuals from species *A*, *B*, and *C* (from left to right) when assuming $K = 2$ or $K = 3$. No case encompasses multiple distinct clustering modes.

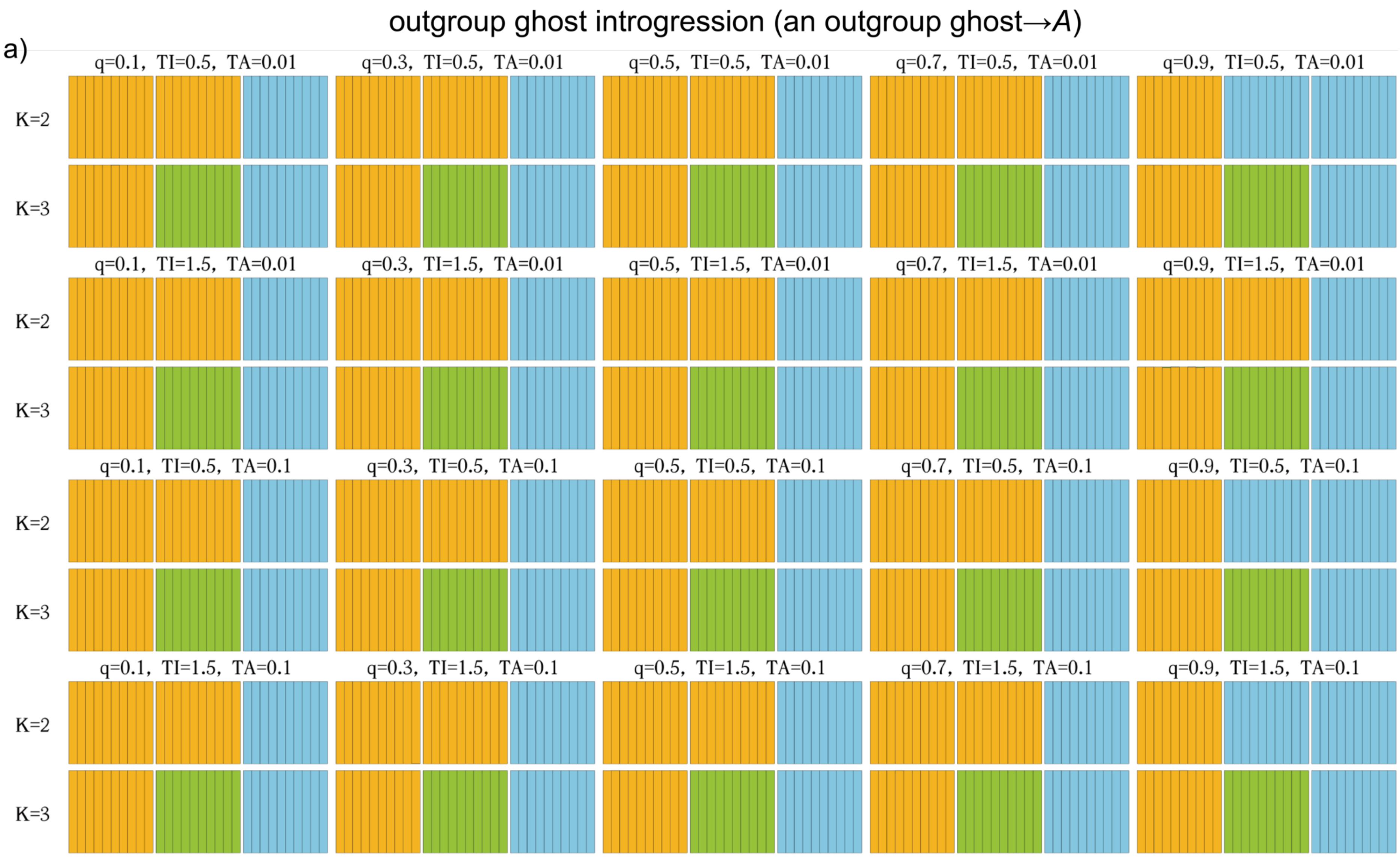
outgroup ghost introgression (an outgroup ghost→A)
a)
q=0.1, TI=0.5, TA=0.01
q=0.3, TI=0.5, TA=0.01
q=0.5, TI=0.5, TA=0.01
q=0.7, TI=0.5, TA=0.01
q=0.9, TI=0.5, TA=0.01
q=0.1, TI=1.5, TA=0.01
q=0.3, TI=1.5, TA=0.01
q=0.5, TI=1.5, TA=0.01
q=0.7, TI=1.5, TA=0.01
q=0.9, TI=1.5, TA=0.01
q=0.1, TI=0.5, TA=0.1
q=0.3, TI=0.5, TA=0.1
q=0.5, TI=0.5, TA=0.1
q=0.7, TI=0.5, TA=0.1
q=0.9, TI=0.5, TA=0.1
q=0.1, TI=1.5, TA=0.1
q=0.3, TI=1.5, TA=0.1
q=0.5, TI=1.5, TA=0.1
q=0.7, TI=1.5, TA=0.1
q=0.9, TI=1.5, TA=0.1
K=2
K=3

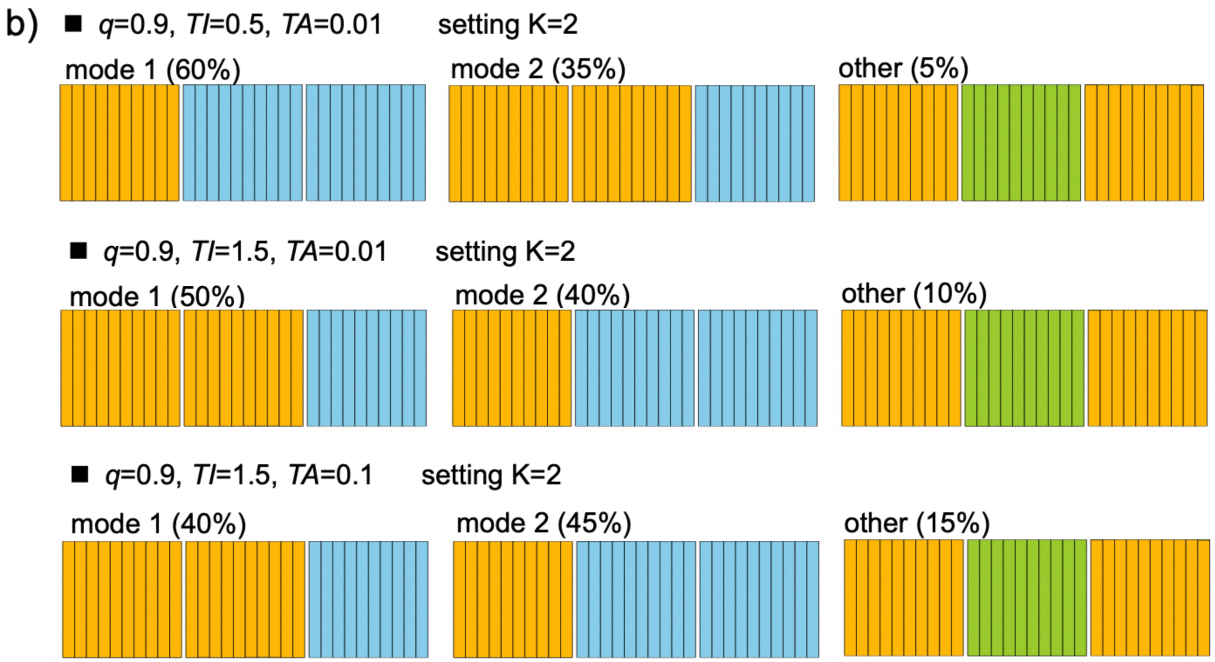


Figure S10. Inferred STRUCTURE clustering plots for $K = 2$ and $K = 3$ under various combinations of parameters in the scenarios of outgroup ghost introgression (Scenario C1 in Fig. 2a). a) The major clustering mode for individuals from species *A*, *B*, and *C* (from left to right). The top of each plot indicates the proportion of genetic material in hybrids inherited from parent *C* ($q$), the time interval between two speciation events (*TI*), and the timing of admixture (*TA*). b) Cases of multiple distinct clustering modes, each supported by at least 20% of the replicates.

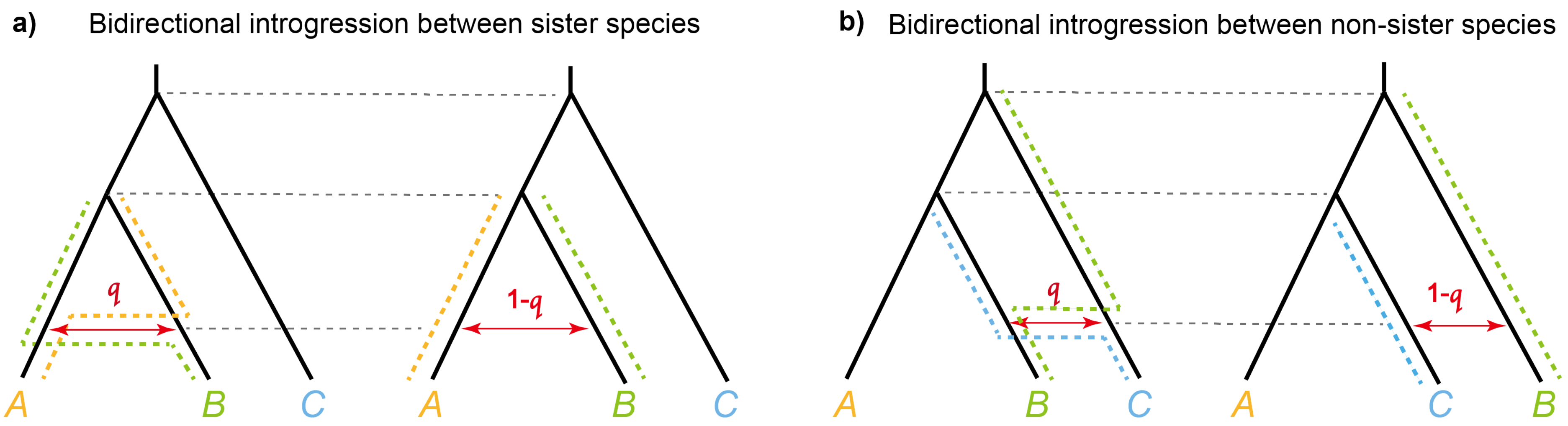


Figure S11. a) A pair of unidentifiable models with a bidirectional introgression event between non-sister species. The first model depicts a species tree *AB*|*C* with a probability of *q* for bidirectional introgression between non-sister species, while the second model illustrates a species tree *AC*|*B* with a probability of 1-*q* for bidirectional introgression between non-sister species. Under the assumption of constant population size, these two models are indistinguishable. b) Unidentifiable parameters for the introgression probability *q* and 1-*q* within the model of bidirectional introgression event between sister species, assuming a constant population size. The dotted lines indicate the main routes taken by sequences sampled from species involved in introgression if the introgression probability in both directions (*q*) exceeds 0.5. For a more in-depth exploration of theoretical proofs, refer to the study by Yang and Flouri (2022).

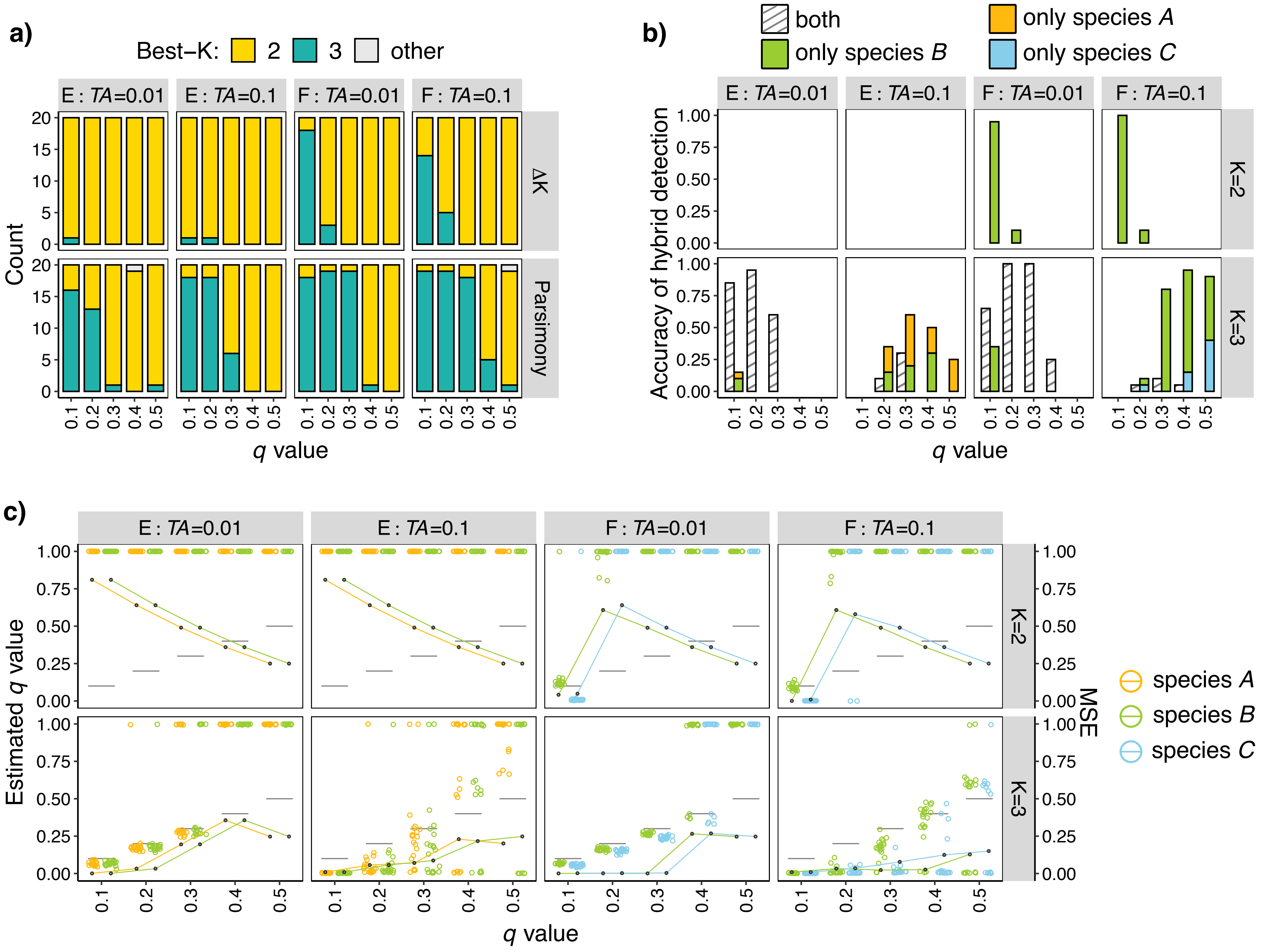

a)
Best−K: 2 3 other
E : TA=0.01
E : TA=0.1
F : TA=0.01
F : TA=0.1
ΔK
Parsimony
Count
q value
b)
both
only species B
only species A
only species C
K=2
K=3
Accuracy of hybrid detection
q value
c)
Estimated q value
MSE
species A
species B
species C
q value

Figure 12. Results for bidirectional introgression scenarios with *TI* = 1.5 (Fig. 3a). a) Best-*K* inference using Δ*K* and parsimony methods. Colored bars represent the numbers of inferences for best-*K* = 2 and best-*K* = 3 among 20 replicates. The strip at the top of each plot indicates the simulated scenarios and the parameter *TA*, while the strip on the right of each plot represents the methods employed. The x-axis indicates the values of *q*. b-c) Plots of results for hybrid detection and admixture proportion estimation assuming *K* = 2 or *K* = 3. The strips located at the right of each plot represent the chosen number of clusters *K*. The x-axis indicates the values of *q*. Results for different hybrid species are labeled in different colors. c) Accuracy of hybrid detection. The y-axis shows the proportion of times that hybrid lineages are successfully identified. The legend categorizes detection results into four distinct types: 'both'—concurrent identification of two hybrid lineages; 'only species *A*', 'only species *B*' and 'only species *C*'—a single hybrid lineage is identified, corresponding to either species *A*, *B*, or *C*, respectively. d) Estimation of admixture proportion *q*-values. Colored points represent the average estimates of *q*-values for hybrid individuals from each hybrid lineage across the datasets, horizontally jittered to avoid clutter. Black lines indicate the true *q*-values, and colored lines with black points indicate MSE values.

a)

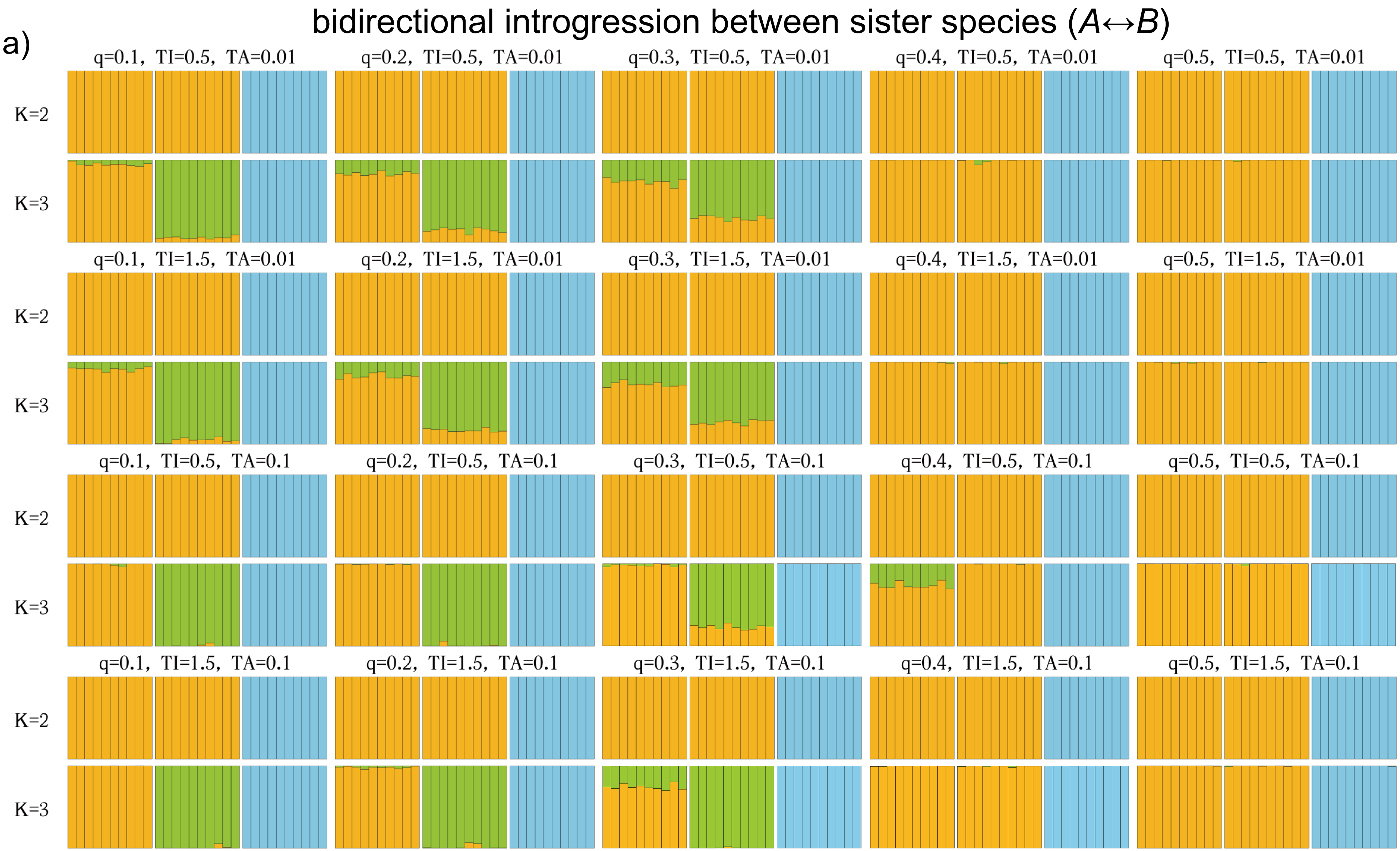

bidirectional introgression between sister species (A↔B)
q=0.1, TI=0.5, TA=0.01
q=0.2, TI=0.5, TA=0.01
q=0.3, TI=0.5, TA=0.01
q=0.4, TI=0.5, TA=0.01
q=0.5, TI=0.5, TA=0.01
K=2
K=3
q=0.1, TI=1.5, TA=0.01
q=0.2, TI=1.5, TA=0.01
q=0.3, TI=1.5, TA=0.01
q=0.4, TI=1.5, TA=0.01
q=0.5, TI=1.5, TA=0.01
q=0.1, TI=0.5, TA=0.1
q=0.2, TI=0.5, TA=0.1
q=0.3, TI=0.5, TA=0.1
q=0.4, TI=0.5, TA=0.1
q=0.5, TI=0.5, TA=0.1
q=0.1, TI=1.5, TA=0.1
q=0.2, TI=1.5, TA=0.1
q=0.3, TI=1.5, TA=0.1
q=0.4, TI=1.5, TA=0.1
q=0.5, TI=1.5, TA=0.1


86

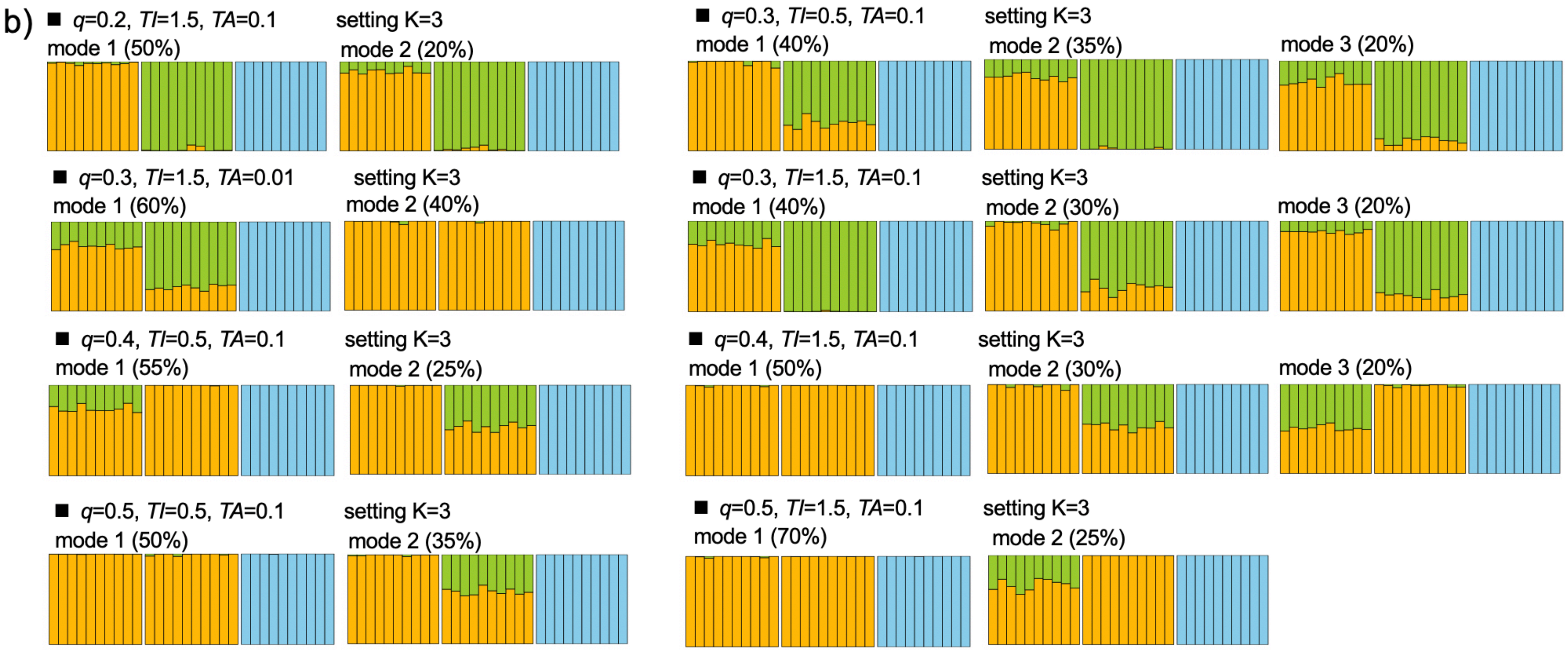


Figure S13. Inferred STRUCTURE clustering plots for $K = 2$ and $K = 3$ under various combinations of parameters in the scenarios of bidirectional introgression between sister species (Scenario E in Fig. 3a). a) The major clustering mode for individuals from species *A*, *B*, and *C* (from left to right). The top of each plot indicates the proportion of genetic material in hybrids inherited from parent *C* (*q*), the time interval between two speciation events (*TI*), and the timing of admixture (*TA*). b) Cases of multiple distinct clustering modes, each supported by at least 20% of the replicates.

bidirectional introgression between non-sister species (*B*↔*C*)

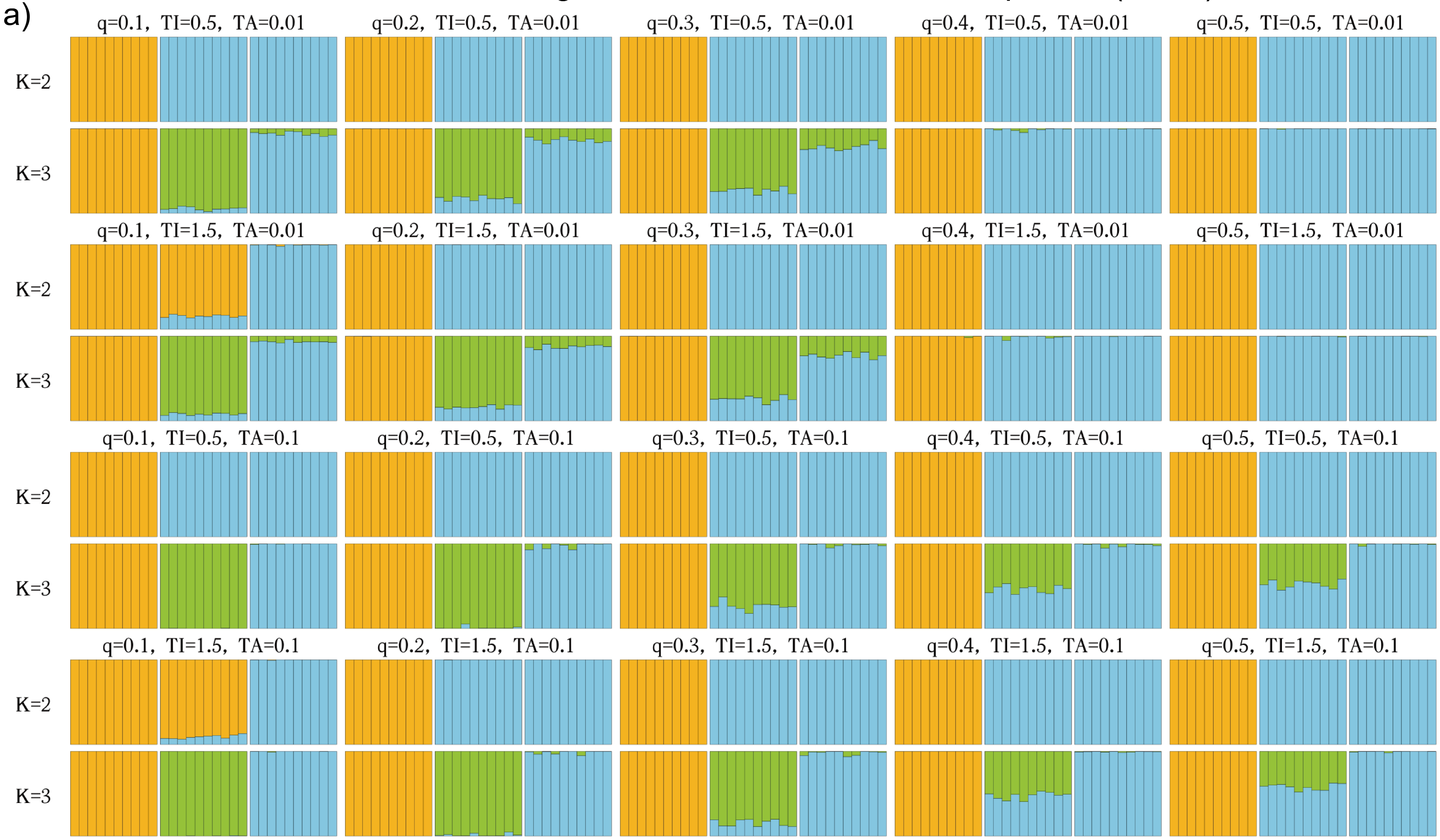


93

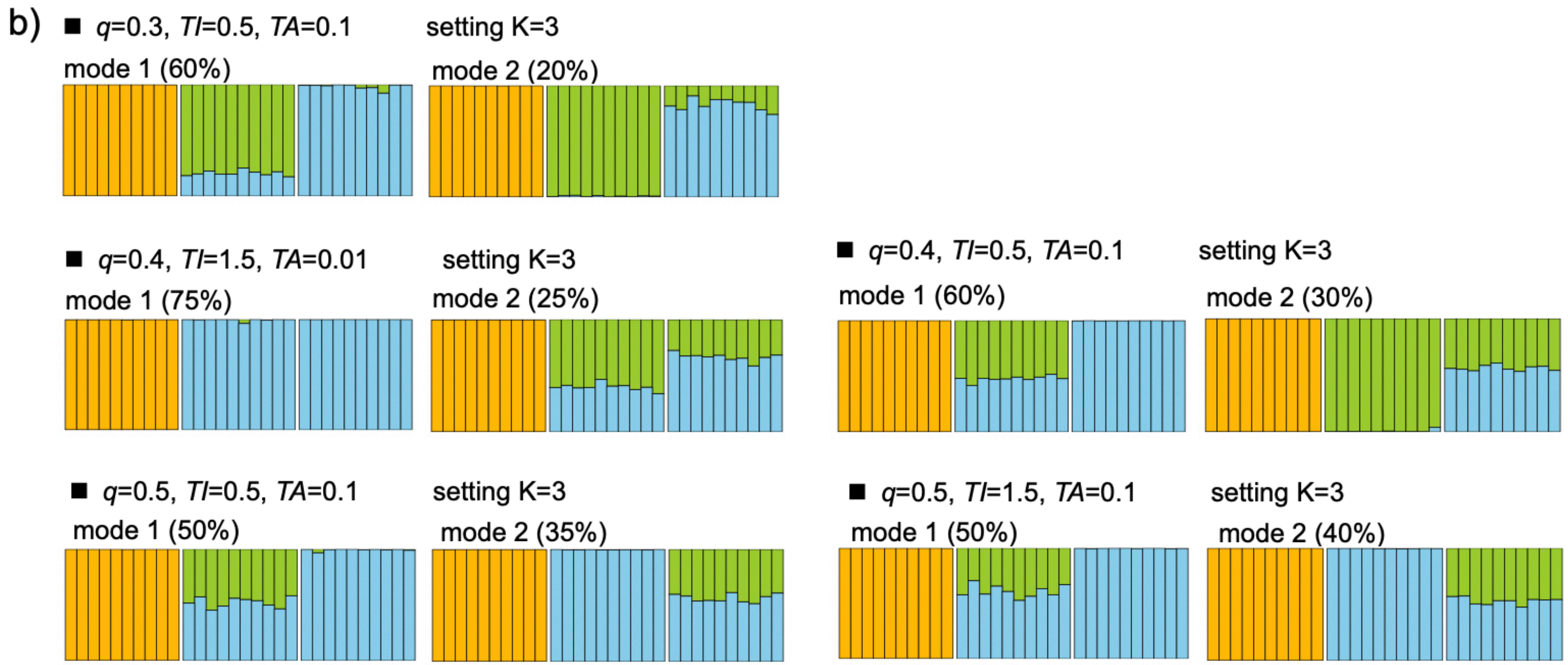


Figure S14. Inferred STRUCTURE clustering plots for $K = 2$ and $K = 3$ under various combinations of parameters in the scenarios of bidirectional introgression between non-sister species (Scenario F in Fig. 3a). a) The major clustering mode for individuals from species *A*, *B*, and *C* (from left to right). The top of each plot indicates the proportion of genetic material in hybrids inherited from parent *C* (*q*), the time interval between two speciation events (*TI*), and the timing of admixture (*TA*). b) Cases of multiple distinct clustering modes, each supported by at least 20% of the replicates.

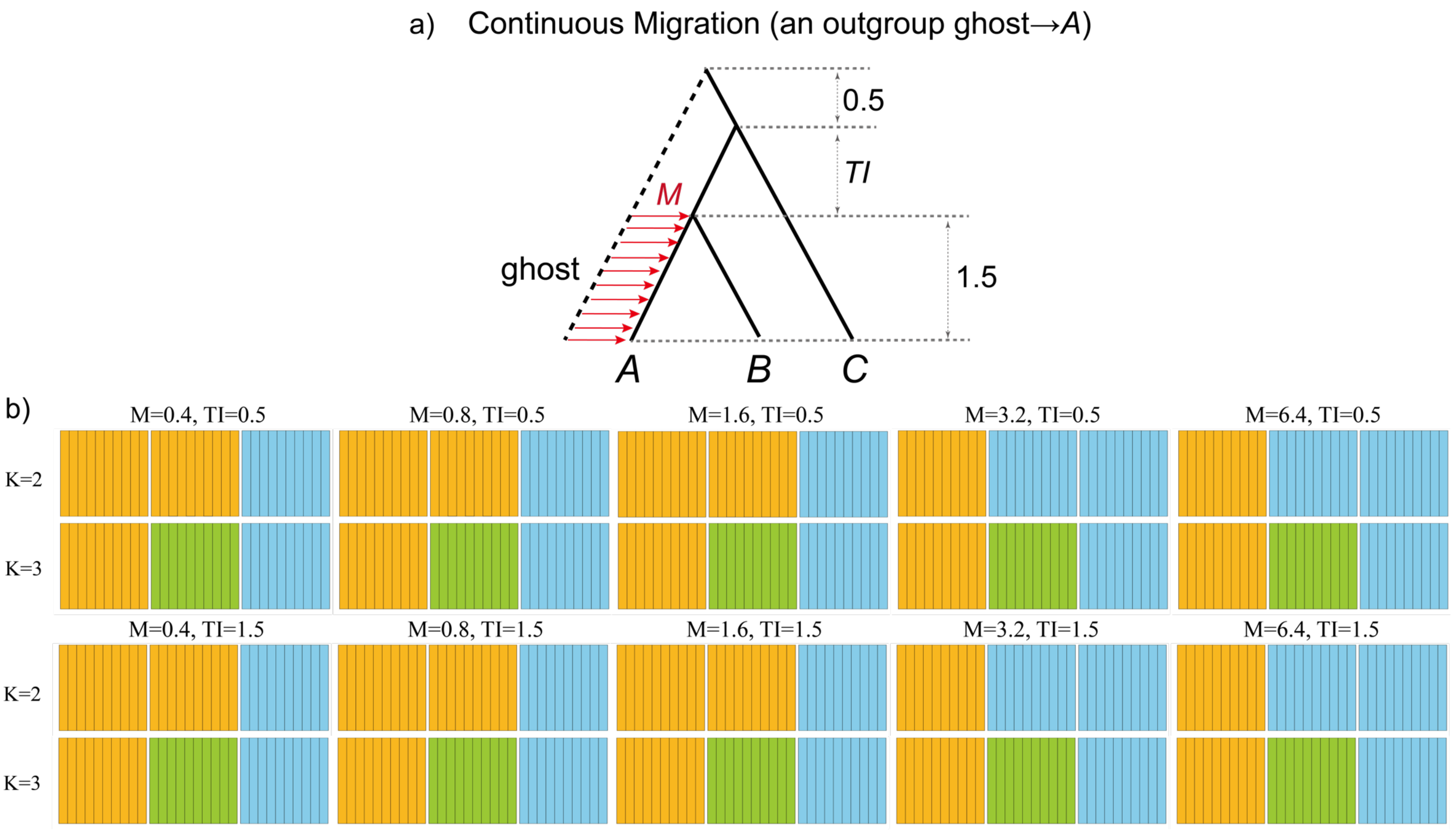
a) Continuous Migration (an outgroup ghost→A)
0.5
TI
M
ghost
1.5
A
B
C
b)
M=0.4, TI=0.5
M=0.8, TI=0.5
M=1.6, TI=0.5
M=3.2, TI=0.5
M=6.4, TI=0.5
K=2
K=3
M=0.4, TI=1.5
M=0.8, TI=1.5
M=1.6, TI=1.5
M=3.2, TI=1.5
M=6.4, TI=1.5
K=2
K=3

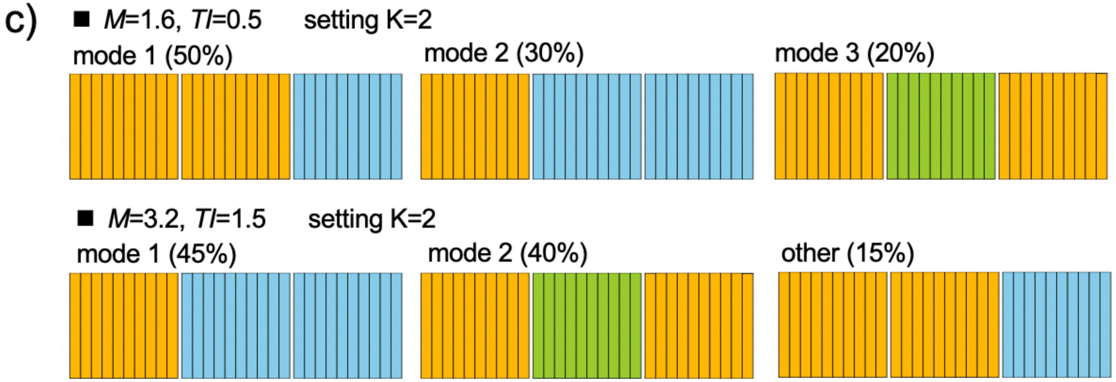


Figure S15. The simulation results of outgroup ghost introgression under continuous migration. a) The simulated scenarios with corresponding parameter settings. Gene flow originates from an outgroup ghost to the sister species *A*, spanning from the divergence of sister species to the present. $M = mN$ represents migration rate, where $N$ is effective population size and $m$ is the proportion of immigrants in the recipient species *A*. *TI* represents the time interval between two speciation events. b-c) Inferred STRUCTURE clustering plots for $K = 2$ and $K = 3$ under various combinations of parameters. b) The major clustering mode for individuals from species *A*, *B*, and *C* (from left to right). c) Cases of multiple distinct clustering modes, each supported by at least 20% of the replicates.

### REFERENCES

Yang Z., Flouri T. 2022. Estimation of cross-species introgression rates using genomic data despite model unidentifiability. Mol. Biol. Evol. 39:msac083.
